# Mechanical stress combines with planar polarised patterning during metaphase to orient embryonic epithelial cell divisions

**DOI:** 10.1101/2023.07.12.548728

**Authors:** Guy B. Blanchard, Elena Scarpa, Leila Muresan, Bénédicte Sanson

## Abstract

The orientation of cell division (OCD) in the plane of epithelia drives tissue morphogenesis and relaxes stresses, with errors leading to pathologies. Stress anisotropy, cell elongation and planar polarisation can all contribute to the OCD, but it is unclear how these interact *in vivo*.

In the planar polarised *Drosophila* embryonic ectoderm during axis elongation, planar OCD is highly variable. We show that both a temporary reversal of tissue stress anisotropy and local compression from neighbouring dividing cells re-orient mitotic spindles during metaphase, independently of interphase cell elongation. Isotropic cells align their OCD to the anterior-posterior (AP) embryonic axis, mediated by tissue-wide planar polarised Myosin II, while the spindle of elongated cells is sterically constrained to cell long axes.

Thus AP-patterning ensures that cell division combines with cell rearrangement to extend the body axis, except when strong local stress anisotropy is dissipated by cells dividing according to their elongation.

## INTRODUCTION

The orientation of cell division (OCD) in the plane of epithelia is important for stress relaxation and tissue morphogenesis. Stress relaxation is achieved through cells dividing along their axis of greatest stress. The orientation of greatest tissue level stress has been shown to influence the orientation of the mitotic spindle and hence the OCD (Campinho et al., 2013; Fink et al., 2011; Mao et al., 2013; Wyatt et al., 2015). Similarly, uniaxial compression has been shown to orient the mitotic spindle perpendicular to the orientation of compression (Lazaro-Dieguez et al., 2015; Lisica et al., 2022). Both anisotropic stress and compression result in cell elongation along or close to the axis of greatest relative stress (Nestor-Bergmann et al., 2019) providing a cell shape cue along which the mitotic spindle can orient, though direct sensing of stress anisotropy has been shown in some examples (Hart et al., 2017; Minc et al., 2011).

During embryonic morphogenesis, cell divisions oriented along the anterior-posterior (AP) axis contribute to axis extension. In vertebrates, such divisions are oriented by planar cell polarity (PCP) (Gong et al., 2004). During *Drosophila* germband extension, AP-patterning orients divisions along AP in the posterior-most abdominal region (da Silva and Vincent, 2007). Also, some cells abutting parasegment boundaries (PSBs) divide along AP because of Myosin II enrichment along the PSB, where it attracts and stabilises spindle ends (Scarpa et al., 2018). However, the majority of germband cells divide along the dorso-ventral (DV) axis, as predicted by their interphase long axis (Scarpa et al., 2018), suggesting that a mixture of patterning and mechanics are involved.

A large body of work supports the Hertwig rule that a cell’s interphase long axis predicts its OCD (Gibson et al., 2011; Gray et al., 2004; Hertwig, 1884; Minc et al., 2011; Wyatt et al., 2015). Because most cells round up during mitosis both in culture (Kunda et al., 2008) and *in vivo* (Chanet et al., 2017) to provide enough room to accommodate mitotic spindle assembly and rotation, the Hertwig rule implies that a molecular mechanism must exist for the mitotic cell to “remember” the interphase cell elongation orientation through mitosis. Such a mechanism has been identified in the *Drosophila* notum, involving the interphase accumulation of Mud (NuMa in vertebrates), which binds to astral microtubules during spindle rotation. Mud is located at tri-cellular junctions which are clustered to the long (distal) ends of cells, providing a memory of interphase topology through mitosis (Bosveld et al., 2016). However, Mud/NuMa is not found at tricellular junctions in all epithelia and divisions do not always align with interphase cell shape (Finegan et al., 2019; Scarpa et al., 2018; Wang et al., 2017). For example, the spindle can be re-oriented in metaphase by mechanical cues (Lazaro-Dieguez et al., 2015) or Myosin II perturbation (Scarpa et al., 2018), over-riding or replacing interphase cues. How mechanical and patterning cues achieve a compromise during OCD is unclear.

Focusing on axis extension in the *Drosophila* embryo, we show that cell divisions can be re-oriented both by a transient reversal of the orientation of tissue stress and by compression exerted by neighbouring dividing cells. These two stress anisotropies exert their influence on the OCD by imposing some cell elongation during metaphase, despite mitotic rounding and independently of interphase cell elongation. We find that if cell elongation in metaphase is strong enough, the spindle is too long for the cell short axis and is sterically constrained to align close to the cell long axis. In addition to this mechanical control of OCD, we also uncover a tissue-wide bias for the mitotic spindle to rotate towards the AP body axis, mediated by planar polarised Myosin II and abrogated in an AP-patterning mutant. AP-patterning ensures AP-oriented divisions in isotropic cells, whilst a compromise between mechanics and AP-patterning is reached in elongated cells. In the latter, the spindle is sterically excluded from the cell short axis but also pulled towards AP, resulting in a spindle orientation at the end of metaphase that is towards AP of the cell long axis. This spindle orientation sets the orientation of anaphase extension and the OCD, with the apparent rotation of the cell long axis in anaphase reflecting the extent to which mechanics has been over-ridden by patterning.

## RESULTS

### Using automated cell tracking to follow the time course of epithelial cell divisions in an intact organism

We set out to analyse the cell divisions in a model epithelium, the germband in the early *Drosophila melanogaster* embryo. Nuclei go through 13 cycles of synchronous divisions before cellularisation, controlled by maternal transcripts (Tadros and Lipshitz, 2009). After cellularisation, cycle 14 to 16 divisions are only locally synchronous and controlled by the zygotic expression of *string*/CDC25 (Edgar and O’Farrell, 1990; Foe, 1989)

To follow cycle 14 epithelial cell divisions in real time, we live imaged the ventral side of the embryo in the trunk/abdominal region (Fig. 1a) for up to 2 hours at 21°C. We acquired two types of dataset, five *shotgun::GFP; sqh::mCherry* at 30 sec. frame intervals for E-Cadherin and Myosin II quantification, which we will call Cad/Myo, and four *ubi-E-Cadherin::GFP;Jupiter::mCherry* at 20 sec frame intervals for quantifying microtubule (MT) spindle movement, which we will call Cad/MT (see Methods). Cell contours at the level of adherens junctions (AJs) using the E-Cadherin signal (Fig. 1b-d and Fig. 1Sa-c) were tracked through time as previously described (Blanchard et al., 2009; Butler et al., 2009; Tetley et al., 2016) (see Methods).

**Figure 1.**
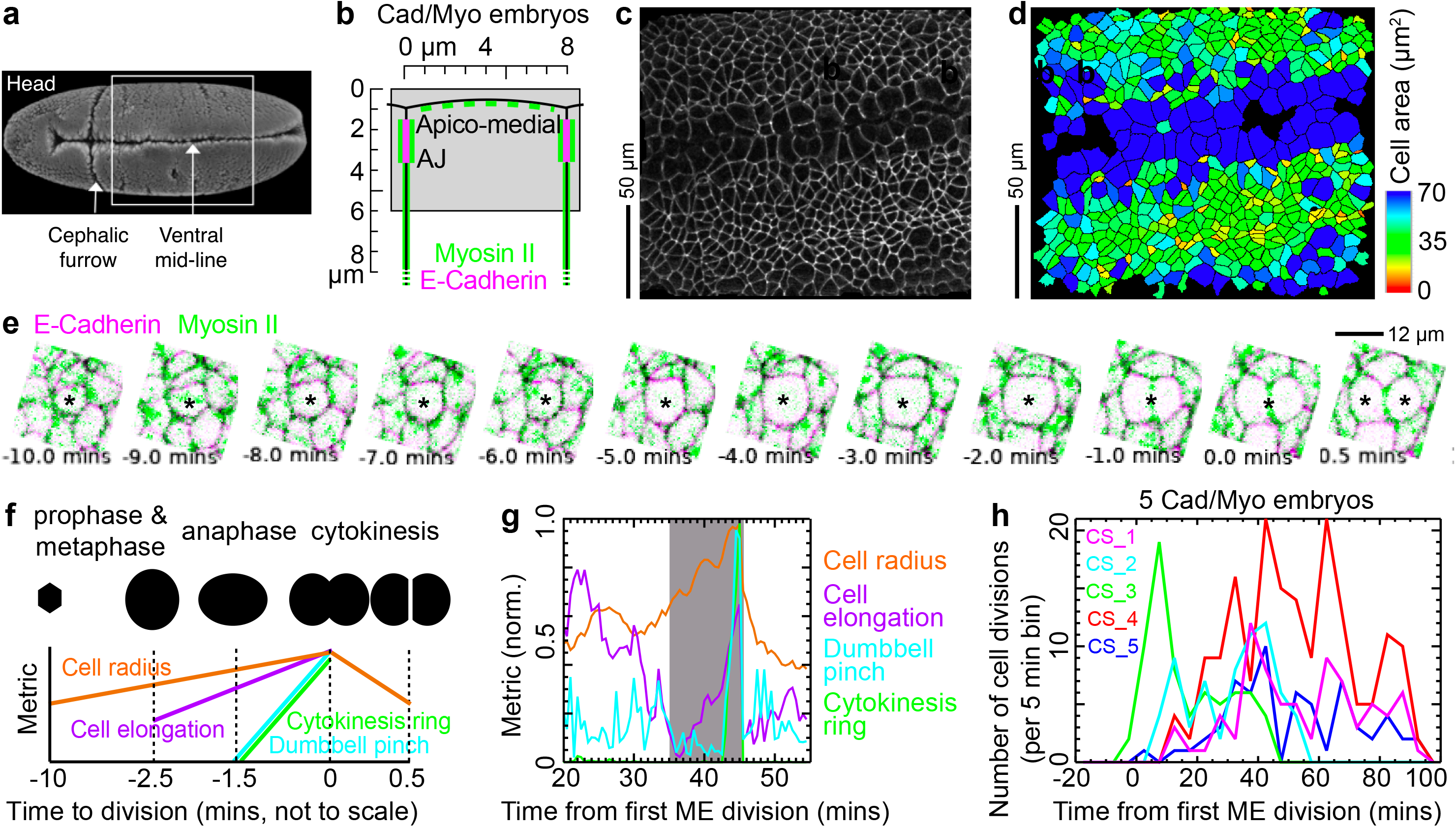
Tracking cell divisions *in vivo*. **a)** Ventral embryo view showing approximate imaged trunk/abdominal region (box). **b)** Schematic of epithelial cell apex showing the depth projection range (grey box) relative to cell apices used to track cells at the level of AJs in Cad/Myo movies. **c)** Projection of E-Cadherin confocal channel at the level of AJs at 0 mins from movie CS_3. Central horizontal band of large cells are mesectoderm (ME) cells, straddling the ventral mid-line. The rest are ectoderm cells. Anterior left. **d)** Segmented cell shapes in (**c**), colour-coded by surface area. **e)** Example of a dividing cell (*) from a Cad/Myo (magenta/green respectively) embryo (see Supplementary Movie 2), from −10 mins to +30 sec after cytokinesis. The cell has been rotated so that orientation of division is horizontal. **f)** Gradients of five metrics are used to classify cytokinesis events. **g)** Example classification of a cytokinesis event, here at 45 mins from first ME division, using metrics color-coded as in (f). Grey box shows the −10 to +0.5 mins window in which mitosis was identified. **h)** Frequency of cell division events over developmental time for five Cad/Myo movies (see Fig. 1Sh For Cad/MT movies). Bin size: 5 mins.

All movies were synchronised to the timepoint in which the first mesectoderm (ME) cell in the field of view reached cytokinesis. This was set to developmental time zero for this study, equivalent to approximately 35 mins after the start of germband extension (GBE) (Tetley et al., 2016). Together, the five Cad/Myo movies span a period from −20 mins to 100 mins (Fig. 1Sd) covering stages 8 to 10 (Bate and Martinez Arias, 1993), with the four Cad/MT movies spanning −10 to 65 mins (Fig. 1Se).

We next developed a classifier algorithm, using an approach similar to (Wang et al., 2017), to identify cell divisions within the population of tracked cells using gradients over time of cell shape and fluorescence metrics (Fig. 1e-g, Fig. 1Sf, g, Supplementary Movie 1 and 2, and see Methods). Most dividing cells in the embryo movies were identified successfully (see Methods), yielding 734 divisions in total over the 9 movies tracked (Fig. 1h and Fig. 1Sh) with a minimum of 10 mins of the mother cell tracked prior to cytokinesis. Having extracted this dataset of cycle 14 divisions during *Drosophila* development, we then assigned these to known mitotic domains.

### Assigning divisions to known mitotic domains

Epithelial cell divisions in the germband occur in mitotic domains which have been identified previously in immunostainings of fixed embryos (Foe, 1989; Hartenstein et al., 1994). We defined two sets of spatial coordinates to assign the cell divisions we identified in real time to specific mitotic domains.

The first identifies a location along the dorso-ventral (DV) embryonic axis for each dividing cell. Tissue converges in DV until 15 mins after the first ME division, after which time the lateral border of the neurectoderm domain is reliably close to 66 µm from the ventral mid-line (VML) across movies (Fig. 2Sa-c and Supplementary Movie 3). We therefore assigned each cell a DV coordinate at 15 mins. We then assigned cells to one of five DV regions that had distinct patterns of cell area over time (Fig. 2a and Fig. 2Sd). These patterns reflected cell division timings and were consistently at the same DV locations across movies (Fig. 2Sc). From ventral to dorsal we named these domains mesectoderm (ME), ventral (VNE), intermediate (INE) and lateral neurectoderm (LNE), and non-neural ectoderm (NNE). Mesectoderm cells abut the ventral midline. Ventral, intermediate and lateral neurectoderm correspond approximately to the expression domains of the three homeodomain transcription factors Vnd (ventral nervous system defective), Ind (intermediate neuroblast defective), Msh (muscle segment homeobox) which pattern the neuroectoderm along DV (Cornell and Von Ohlen, 2000). The non-neural ectoderm extends between the LNE and the out of view amnioserosa.

**Figure 2.**
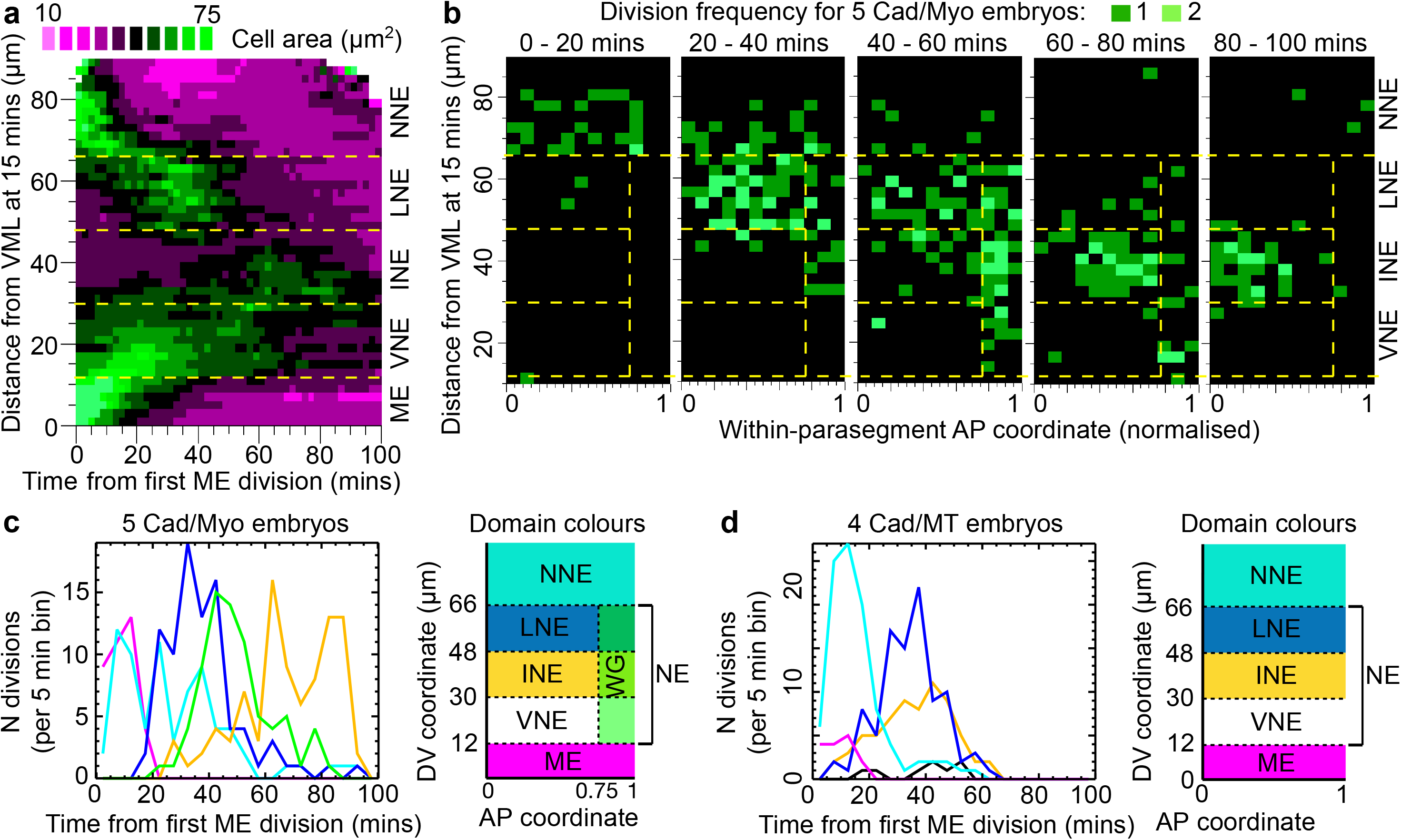
Assigning divisions to known mitotic domains. **a)** Average cell areas for pooled data from five Cad/Myo embryos. Yellow dashed lines show manually assigned borders between DV domains with different patterns of cell shape. The DV y-axis is a co-moving coordinate frame, assigned to cells at 15 mins. Cell area increase is one of the cell division identifiers. **b)** Frequency of cell division events from five Cad/Myo embryos, broken down by DV co-moving coordinate, AP within-parasegment coordinate and 20-min development time bins. Yellow dashed grid shows imposed AP and DV boundaries. ME cells are not plotted because they are not assigned within-parasegment coordinates. All are cycle 14 divisions and no cells divided twice. **c)** Frequency of cell divisions identified over time in Cad/Myo embryos in the five spatio-temporal domains identified in (b). Bin size is 5 mins. **d)** Frequency of cell divisions identified over time in Cad/MT embryos in four spatio-temporal domains. Bin size is 5 mins.

The second set of spatial coordinates, along the AP embryonic axis, identifies the relative location of each dividing cell within each parasegment. In Cad/Myo embryos, parasegment borders are identified by the presence of an actomyosin cable (Monier et al., 2010) (Fig. 2Se, f) between which a within-parasegment AP coordinate (normalised between 0 and 1) is assigned to each cell, as previously (Fig. 2Sg and Supplementary Movie 3) (Tetley et al., 2016).

Across five Cad/Myo embryo movies we consistently identified 6 distinct cell division domains in our spatio-temporal mapping (Fig. 2b, c and Fig. 2Sh). ME cells (less than 12 µm from VML) divide at the same time (0-20 mins) as the most lateral NNE domain (> 66 µm from VML). Next, the LNE domain (48-66 µm, 20-40 mins) divides before the INE domain (30-48 µm, 60-100 mins), the latter cells dividing in an anterior-ward wave. Most VNE cells (12-30 µm) divide after 100 mins, beyond the end of our quantification. Within parasegments, the anterior three-quarters behave as above, but the posterior quarter, containing wg-expressing cells, behaves uniquely. This WG domain contains cells from LNE, INE and VNE domains dividing together (40-60 mins). The temporal sequence of divisions we observed in the domains ME, NNE, LNE, WG, INE, VNE agree with the temporal sequence of equivalent domains previously identified from fixed samples (Foe, 1989). In Cad/MT embryos we could not identify the WG domain because there was no Myosin II channel (Fig. 2Si and Fig. 2d), so for analyses involving Cad/MT embryo data, we only distinguished between the five DV cell type domains.

### The time course of mitosis is similar in different mitotic domains

To compare cell divisions, we first asked whether there are differences in the rate of progress through mitosis between domains, such as the start of rounding, appearance of the mitotic spindle and cytokinesis ring. For both Cad/Myo and Cad/MT movies, we synchronised each cell division time course relative to the last movie frame before a new E-Cadherin junction forms between new daughter cells (Fig. 1e), defining a time zero for all cell divisions (Supp Movie 1 and Fig. 3Sa).

Aligning all 734 divisions relative to this time zero, we find that the evolution of average cell shapes is remarkably similar across all mitotic domains (Fig. 3a for Cad/Myo and Fig. 3Sb for Cad/MT). Cell area increase in the AJ plane is first detectable at −12 mins, continuing through to cytokinesis. ME cells are apically larger than the more columnar cells in the neurectoderm domains, but they follow a similar time-course of area increase. Cell elongation starts to decrease at −11 mins, continuing until −2.5 mins, and then increasing again during anaphase elongation (black line in Fig. 3a and Fig. 3Sb). Apicomedial Myosin II (Fig. 1b) density starts to reduce at −13 mins (Fig. 3b), the earliest detectable change that we could find associated with cell division. Cytoplasmic MT fluorescence density (Fig. 3Sc, d), evidence of the mitotic spindle, peaks at −3 mins just before the start of anaphase (Fig. 3b).

**Figure 3.**
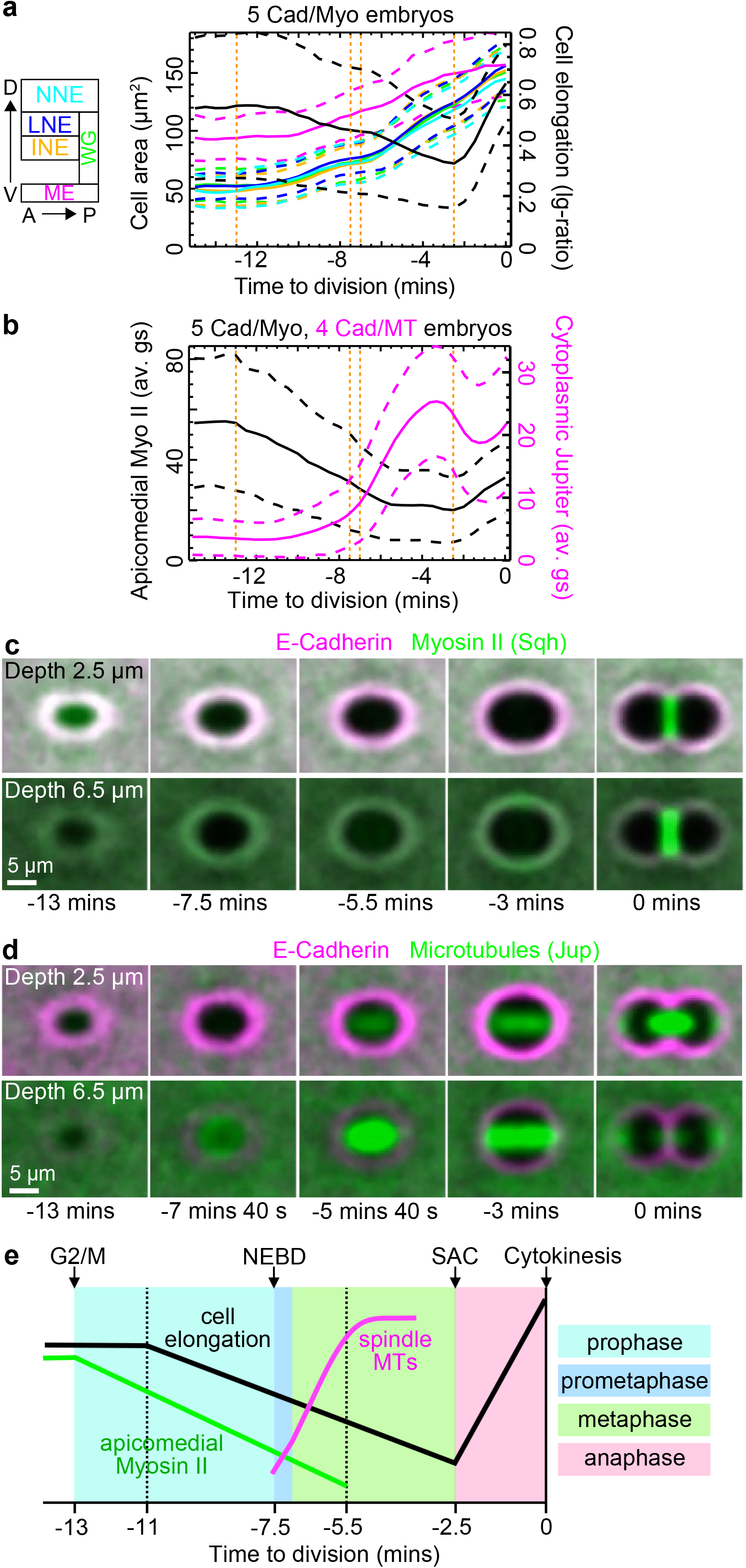
The time course of mitosis. **a)** Cell shape through mitosis at the level of AJs for Cad/Myo embryos. Left y-axis, cell area for five mitotic domains, colour-coded according to parasegment schematic. Right y-axis, cell elongation ratio (black line). **b)** Fluorescence patterns through mitosis. Left y-axis, apicomedial Myosin II density for Cad/Myo embryos (black line). Right y-axis, cytoplasmic MT density for Cad/MT embryos (magenta line). Orange dotted lines in (a) and (b) show detectable start of mitosis at −13 mins, NEBD at −7.5 mins, start of metaphase −7 mins and start of anaphase −2.5 mins. Pooled cell means are shown +/−SD. **c)** Average cell fluorescence around tracked cell centroids from Cad/Myo embryos at depths 2.5 µm (top) and 6.5 µm (bottom) from the surface of the epithelium. Data pooled from 411 ectoderm cells. **d)** Average cell fluorescence around cell centroids from Cad/MT embryos at depths 2.5 µm (top) and 6.5 µm (bottom) from the surface of the epithelium. Data pooled from 270 ectoderm cells. OCD is set to horizontal for all cells in (c) and (d). **e)** Schematic summary of the time course of *Drosophila* round 14 divisions. G2/M; effective G2 to mitosis transition. SAC; spindle assembly checkpoint marking anaphase onset (see Methods).

For a visual confirmation of mitotic timings, we generated a representation of average cell fluorescence through division (see Methods). We assigned to each cell a division axis, the orientation between daughter centroids in the first frame after division, which allowed us to co-orient dividing cell behaviour leading up to cytokinesis (Fig. 1e and Fig. 3Se). We then overlaid and averaged fluorescence intensities surrounding the cell centroids of all divisions at the level of AJs and 4 µm deeper at a sub-AJ level for Cad/Myo (Fig. 3c and Supplementary Movie 4) and Cad/MT (Fig. 3d and Supplementary Movie 5) movies. In the Jupiter channel, the low fluorescence signal of MTs at −7’40” is from centrosomes at the base of the nucleus just prior to nuclear envelope breakdown (NEBD). The mitotic spindle, which appears bright, is then variably oriented in the plane at −5’40” then oriented along the future OCD at −3 mins. Given that the cytokinesis ring in the Myosin II channel at 0 mins is not yet visible at −3 mins, this is still metaphase.

Putting the above information in sequence (Fig. 3e), we define the initial decrease of of apicomedial Myo II at −13 mins as our effective G2/M transition. Prometaphase starts at - 7.5 mins with NEBD, followed immediately by the formation of the metaphase spindle. Metaphase then spans −7 to −2.5 mins, when anaphase elongation commences. This tight mitotic temporal schedule is adhered to on average by all mitotic domains. We next ask how the orientation of cell division (OCD) varies and whether it differs between mitotic domains.

### Metaphase cell shape tracks tissue tension

We set out to investigate how the OCD is distributed in the mitotic domains we identified. Overall, we observed that the orientation of cell division is biased towards DV (Fig. 4a), which confirms our previous finding by immunostaining of fixed embryos (Scarpa et al., 2018). However, the distribution of angles is widely spread from AP to DV. We noticed that the early cell division domains (NNE, ME) were AP-biased, while the later NE domains were DV-biased (Fig. 4a, Fig. 4S1a). Pooling domains into these two groupings, the average division orientation changes from AP- to DV-oriented around 20 mins (Fig. 4b).

**Figure 4.**
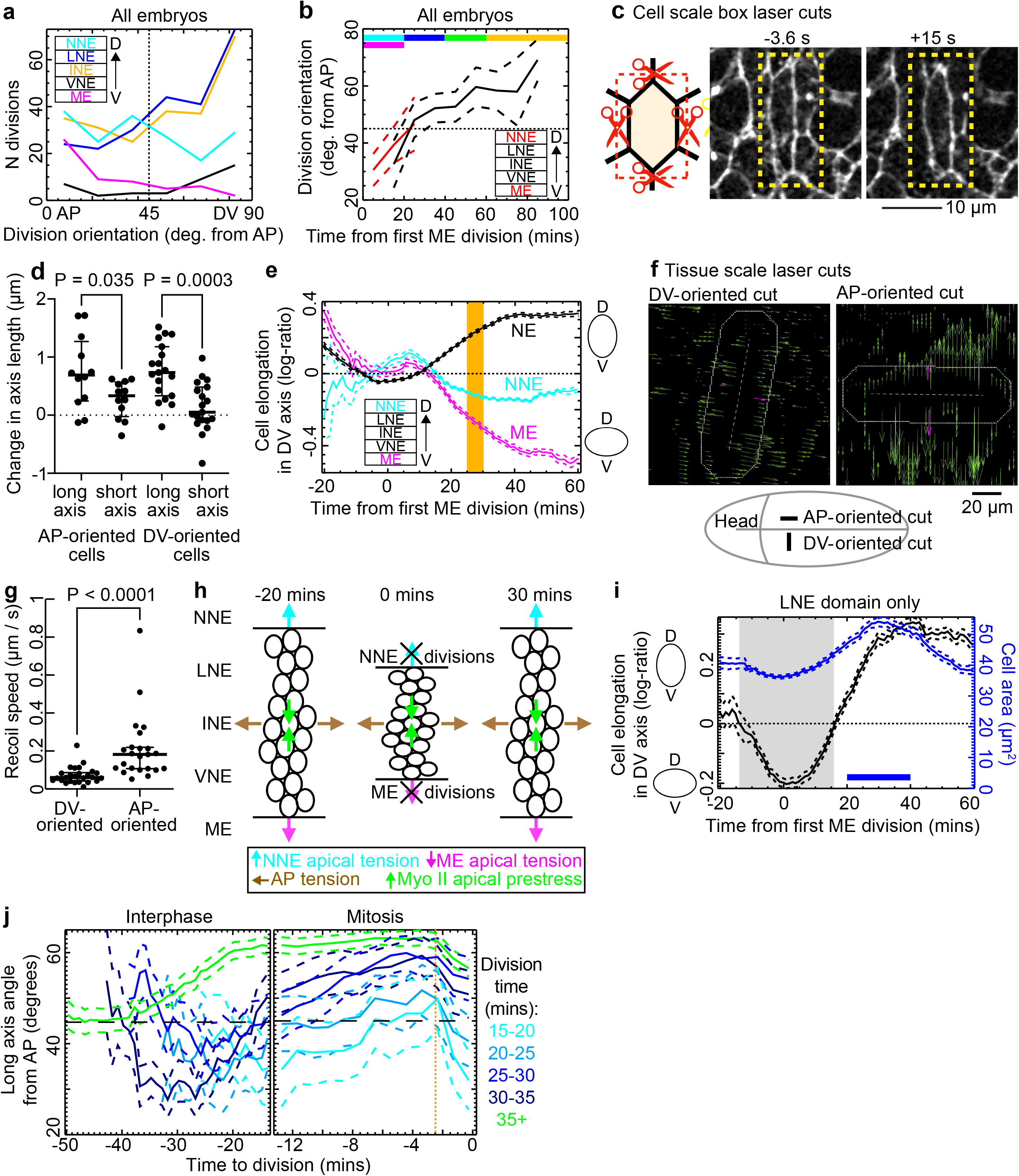
Metaphase cell shape tracks tissue tension. **a)** Frequency of the OCD by mitotic domain. Bin size is 15 degrees. **b)** OCD over developmental time for ME and NNE (red) and NE (black) cells. Horizontal bars show epochs of division for each mitotic domain, coloured as in (a). **c)** For box laser cut ablations, schematic shows how a rectangular template is constructed, cutting neighbour-neighbour junctions around the focal cell. Stills from a box ablation movie, just before and 15 s after the cut, showing the retraction of the focal cell and severed neighbour-neighbour junctions. Dashed yellow box shows cut. **d)** Reduction in axis lengths after box ablation broken down by orientation of cell long axis before ablation (Paired t-tests; AP-oriented, N = 12; DV-oriented, N = 19) (see Fig. 4S1b-e). **e)** Cell elongation in the DV axis for ME, NNE and pooled NE domains over developmental time. Tissue laser cuts were performed at 25-30 mins (orange bar). **f)** AP (left) and DV (right) component of the PIV flow field in Sqh-GFP channel 2.9 sec after tissue scale laser cuts. Cut sites and orientation are indicated by the dashed line. Schematic shows the location of AP- and DV-oriented tissue scale cuts in the ventral extended germband. **g)** Initial retraction speeds after tissue-scale laser ablation (Two sample t-test; N = 25 each). **h)** Model for the temporary reversal of the orientation of greatest tissue stress. Divisions in the neighbouring ME and NNE tissues around 0 mins temporarily release tension in (maybe compress, see Fig. 7) the NE in DV, resulting in AP tension temporarily overtaking DV tension. **i)** Cell elongation in the DV axis and cell area over developmental time for the LNE mitotic domain only, which divides between 20-40 mins (horizontal blue bar). The stress inversion, defined as where cells are AP-elongated, is shown in grey. **j)** Orientation of NE cells in interphase and mitosis (note different x-axis scales), separated into 5-min division time bins. Early dividing LNE cells can be seen changing their long axis from AP to DV-oriented from interphase to anaphase, as the tissue stress changes from AP- to DV-dominated (see (i)). The green ‘35+’ min bin includes all NE cells that divide between 35 and 100 mins. Dotted orange line shows start of anaphase.

We hypothesised that this change could be due to a change in the orientation of greatest tissue stress. During axis extension, the germband is subject to an extrinsic AP-oriented pull (Butler et al., 2009; Collinet et al., 2015; Lye et al., 2015) and to intrinsic DV-oriented contractility driven by polarised Myosin II at cell-cell junctions (Rauzi et al., 2008; Tetley et al., 2016; Zallen and Wieschaus, 2004). Tension is greater in DV during the fast phase of GBE (Collinet et al., 2015), after which stress is likely to be more balanced or AP-oriented during ME divisions (Camuglia et al., 2022; Wang et al., 2017). It is not known what happens to the orientation of tissue stress during NE divisions, so we probed the tissue stress anisotropy in three ways:

1) The cell long axis has been shown to be a reliable readout of the axis of greatest local stress (Nestor-Bergmann et al., 2018; Nestor-Bergmann et al., 2019). We tested this in the NE by doing ‘box’ laser cuts to isolate the apical surface of interphase cells from all neighbours (Fig. 4c). Indeed, the main direction of recoil is systematically correlated with the cell long axis, irrespective of the orientation of the cell (Fig. 4d and Fig. 4S1b-e). Thus, embedded within a tissue that has a global anisotropic stress, cell elongation is associated with local anisotropic stress, greater in the cell long axis than short axis. We therefore compared AP and DV cell lengths in the NE as a proxy for the orientation of greatest local stress. NE cell elongation in DV reduced dramatically at the end of the fast phase of GBE to - 10 mins, when the cell long axis flipped from DV to AP (Fig. 4e) as cells reduced area (Fig. 4S1f). This reversed back to DV-elongated at around +10 mins, with cells continuing to elongate in DV to 40 mins. Cell shapes therefore predict a switch in tissue stress anisotropy in the NE to AP-oriented at −10 mins and then back to DV-oriented at +10 mins.
2) We performed tissue scale laser cuts on the apical myosin meshwork in perpendicular orientations at 25-30 mins, after ME and NNE divisions and near the peak of DV cell length (orange bar in Fig. 4e). We found that indeed tissue stress anisotropy has returned to being strongly DV-oriented (Fig. 4f, g and Methods).
3) We wanted to know whether this tissue stress anisotropy reversal is an apical feature transmitted through AJs. We measured the 3D wedge shape and tilt of NE cells in a Cad/Myo embryo by tracking cells at a sub-AJ depth and matching tracked cells to our AJ-level tracking (Fig. 4S2 and see Methods). During the tissue stress reversal (5 mins in Fig. 4e), cells are negatively wedge-shaped (smaller apices) in DV (Fig. 4S2f) and tilted towards each other (Fig. 4S2g). After the tissue stress reversal (45 mins), cells become positively wedge-shaped (stretched apices) in DV and tilted outwards. Thus, the tissue stress reversal is transmitted most strongly at the level of AJs.

This gave us a clue as to what might be causing the reversal in tissue stress. Dividing cells round up predominantly by pulling up their bases, leading to apical expansion. As in other tissues (Gupta et al., 2021) this is likely to release apical tension in, and/or induce compression in, nearby cells (see Fig. 7). Divisions in the ME and NNE, the cell domains on either side of the NE, could therefore be the cause of the loss of DV tension in the NE (Fig. 4h. This is supported by the tight temporal coupling at −5 mins between the metaphase rounding of the ME and NNE cells (Fig. 4S1f) and the apical DV shortening of the NE domain in between (Fig. 4e). Once the ME and NNE have finished dividing at 15-20 mins (Fig. 4S1f and Fig. 2c, d) they progressively contract in DV to 30 mins, stretching the NE tissue and the orientation of greatest stress returns to DV.

Above we have provided evidence that supports a reversal in tissue stress in the NE during ME and NNE divisions. Given that LNE divisions start immediately afterwards (20-40 mins, Fig. 2c, d), we asked how strictly do cell elongation and OCD track changes in tissue stress. Early LNE divisions will be exposed to the switch in tissue stress during mitosis while the later ones will only experience it during interphase. LNE cells switch from AP- to DV-oriented at 15 mins and are maximally stretched in DV by 30 mins (Fig. 4i). Breaking LNE divisions down by 5-min division time bins, we find that dividing cells become progressively more DV-oriented during mitosis in all bins, crossing from AP- to DV-oriented at the time expected from the change in tissue stress (Fig. 4j). Later divisions unaffected by the tissue stress reversal do not change orientation during mitosis (35+ bin). Thus, cell shape during mitosis instantaneously tracks tissue stress on average. Surprisingly, however, this does not continue after −2.5 mins, the onset of anaphase, when cell orientations rotate dramatically to AP (Fig. 4j).

We conclude that there is a switch in NE tissue tension anisotropy during early divisions, likely caused by divisions in adjacent ME and NNE tissues (see Fig. 4h and Discussion). The shapes of dividing cells instantaneously track this switch in tissue tension in mitosis through to metaphase. During anaphase, however, there is a rotation of the long axis orientation towards AP that we investigate next.

### An AP-oriented cue attracts the mitotic spindle independently of tissue stress

The rotation of the cell long axis towards AP during anaphase could either be because cells have a memory of interphase AP orientation, or because there is a stress-independent bias to AP. We note that for divisions between 35 and 100 mins, long after the reversal of tissue stress, there is still an anaphase long axis rotation to AP (Fig. 4j, green line). Indeed, across all our study period the division angle is consistently rotated to AP from the metaphase (−4 mins) long axis orientations (Fig. 5a). This indicates that the anaphase rotation is not linked to the tissue stress reversal and hence not due to an interphase memory of stress.

**Figure 5.**
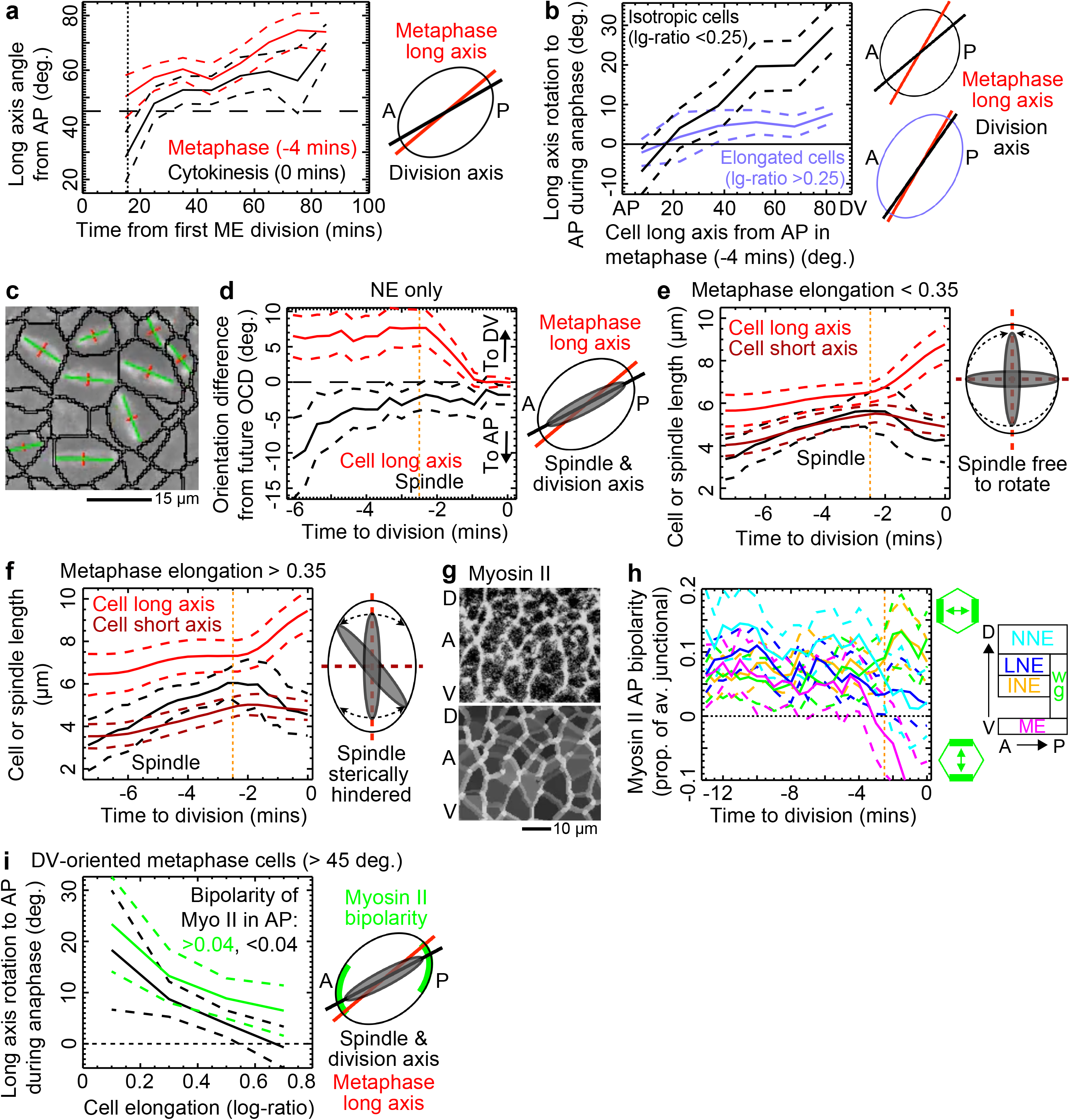
An AP-oriented cue attracts the mitotic spindle independently of mechanics. **a)** Metaphase cell long axis orientation compared to division axis at cytokinesis (OCD) across developmental time. Vertical dotted line marks the end of the tissue stress reversal. **b)** Magnitude of rotation to AP in anaphase (difference between long axis orientations at metaphase and cytokinesis) for isotropic and anisotropic cells. **c)** Example segmented spindles (green lines, principal components; red lines, orthogonal components) drawn from movie of Jupiter tracking (Supplementary Movie 6). **d)** Angular difference, to AP or DV, between both the cell long axis and spindle and the OCD for NE cells. **e-f)** Comparison of the cell long and short axis lengths with spindle length for **e)** isotropic and **f)** elongated cells. **a)** Top; raw junctional Myosin II fluorescence. Bottom; segmented junctions, colour-coded by junctional Myosin II density (greyscale). **b)** Myosin II bipolarity projected onto the AP-axis for different mitotic domains during mitosis. Positive bipolarity is AP-oriented. **c)** Long axis rotation to AP in anaphase for DV-oriented metaphase cells, separated into high (green) and low (black) myosin AP-oriented bipolarity. In all panels, averages are plotted +/−95% CIs (dashed lines). Vertical orange dotted lines mark the start of anaphase. Data pooled from 5 Cad/Myo and 4 Cad/MT embryos in **a-b**, from 4 Cad/MT embryos in **d-f** and from 5 Cad/Myo embryos in **h-i**.

We wanted to understand the magnitude of anaphase rotation, which appeared larger earlier (Fig. 5a, < 30 mins). Separating end of metaphase cells (−4 mins) into either isotropic or elongated, isotropic cells have a much larger rotation towards AP compared to elongated cells, up to 30 degrees for the most DV-oriented of the isotropic cells (Fig. 5b). Elongated cells also rotated significantly to AP but only by around 5 degrees on average. This suggests that the rotation to AP is ubiquitous, though constrained in elongated cells.

To understand whether the mitotic spindle rotates with the cell long axis or whether the long axis rotates to the spindle, we tracked spindles in 3D within each mitotic cell, from their formation after NEBD (−7 mins) until cytokinesis (Fig. 5c, Fig. 5Sa-c, Supplementary Movie 6 and Methods). Strikingly, at the end of metaphase (−2.5 mins), the spindle is more closely aligned to the future OCD than to the cell long axis in NE cells (Fig. 5d). During anaphase, the long axis then rotates towards the spindle axis and towards AP. A similar but more dramatic rotation to AP is seen in the NNE and ME (Fig. 5Sd-e), which we hypothesised could be because cells in these domains are more isotropic (Fig. 5Sf and Fig. 5b). Indeed, comparing spindle lengths with the short axes of metaphase cells, spindles in isotropic cells experience no constraint on their orientation (Fig. 5e) but short axes of elongated cells are significantly shorter than their spindle lengths (Fig. 5f). Steric hindrance of the spindle by the cell short axis therefore explains why elongated cells rotate less towards AP in anaphase than isotropic cells and provides a mechanism for how anisotropic stress, through cell elongation, constrains OCD.

So, what is the cue attracting spindles away from cell long axes and towards AP? Various proteins are planar polarised to (or away from) DV-oriented junctions downstream of AP-patterning (Pare and Zallen, 2020) and could attract (or repel) the spindle. Of these, Myosin II is strongly polarised to DV-oriented junctions during rapid GBE (Rauzi et al., 2008; Tetley et al., 2016; Zallen and Wieschaus, 2004) and is embedded in the actin cortex that has an instructive role in spindle orientation (di Pietro et al., 2016). For example, we have previously shown that cells adjacent to the Myosin II-enriched PSBs orient their divisions towards the PSB, along AP, even if they are mildly elongated in DV (Scarpa et al., 2018). We therefore asked whether Myosin II attracts the spindle towards AP and away from cell long axes, using the OCD as a proxy for metaphase spindle orientation (see above).

We calculated the orientation of Myosin II bipolarity and unipolarity and projected these onto the AP axis (Fig. 5Sg, Fig. 5g and Methods) (Tetley et al., 2016). Both measures are strongest during GBE (−20 mins to 0), as expected, but some polarisation remains past axis extension (0 to 100mins) (Fig. 5Sh, i). Myosin II bipolarity, but not unipolarity, is AP-oriented on average in all domains at the developmental time at which they divide (Fig. 5Sh). The unipolarity signal, though strong at PSBs, on average changes orientation to DV at ∼40 mins due to a progressive strengthening at some AP-oriented junctions (Fig. 5Si). Focusing on mitotic cells only, Myosin II bipolarity is significantly AP-oriented in all mitotic domains through metaphase (Fig. 5h). Furthermore, stronger Myosin II bipolarity is associated with greater long axis rotation to AP (Fig. 5i).

We conclude that Myosin II is AP-oriented in all mitotic cells in the germband epidermis and our correlations suggest that it biases the spindle orientation away from cell long axes and towards AP in all cells. The spindle in elongated cells is sterically hindered by the cell short axis, leading to smaller spindle axis rotations, but in isotropic cells the spindle axis can rotate freely.

### Anaphase long cell axis rotation towards AP is lost in an AP-patterning mutant

Myosin II planar polarization is controlled by AP patterning in the *Drosophila* germband (Bertet et al., 2004; Zallen and Wieschaus, 2004). To test whether Myosin II is responsible for attracting the spindle towards AP, we imaged three double mutant (−/−) and three heterozygote (+/−) *Knirps, Hunchback* (*KniHb*) embryos into which we had introduced Gap43;Sqh (Gap/Myo) fluorescent reporters. *KniHb-/-* embryos lack essential gap genes responsible for setting up AP-patterning in the trunk and abdomen (Irvine and Wieschaus, 1994).

ME divisions in the WT had the greatest anaphase long axis rotation to AP because of their larger and isotropic apices. We therefore focused on these divisions in the *KniHb* embryos. We also captured some LNE divisions, though these were unlike WT in often being asymmetric in the plane, or out of plane. We first checked gross phenotypes, finding that ME divisions in *KniHb*+/−were indistinguishable from WT, being predominantly AP-oriented and resulting in a ME domain 2-3 cells wide after divisions (Fig. 6a and Fig. 6Sa) and with clear PSBs in the Myosin II channel. By contrast, *KniHb*-/-ME divisions were variably oriented, resulting in a 4-5 cell wide domain after division, and with no PSBs.

**Figure 6.**
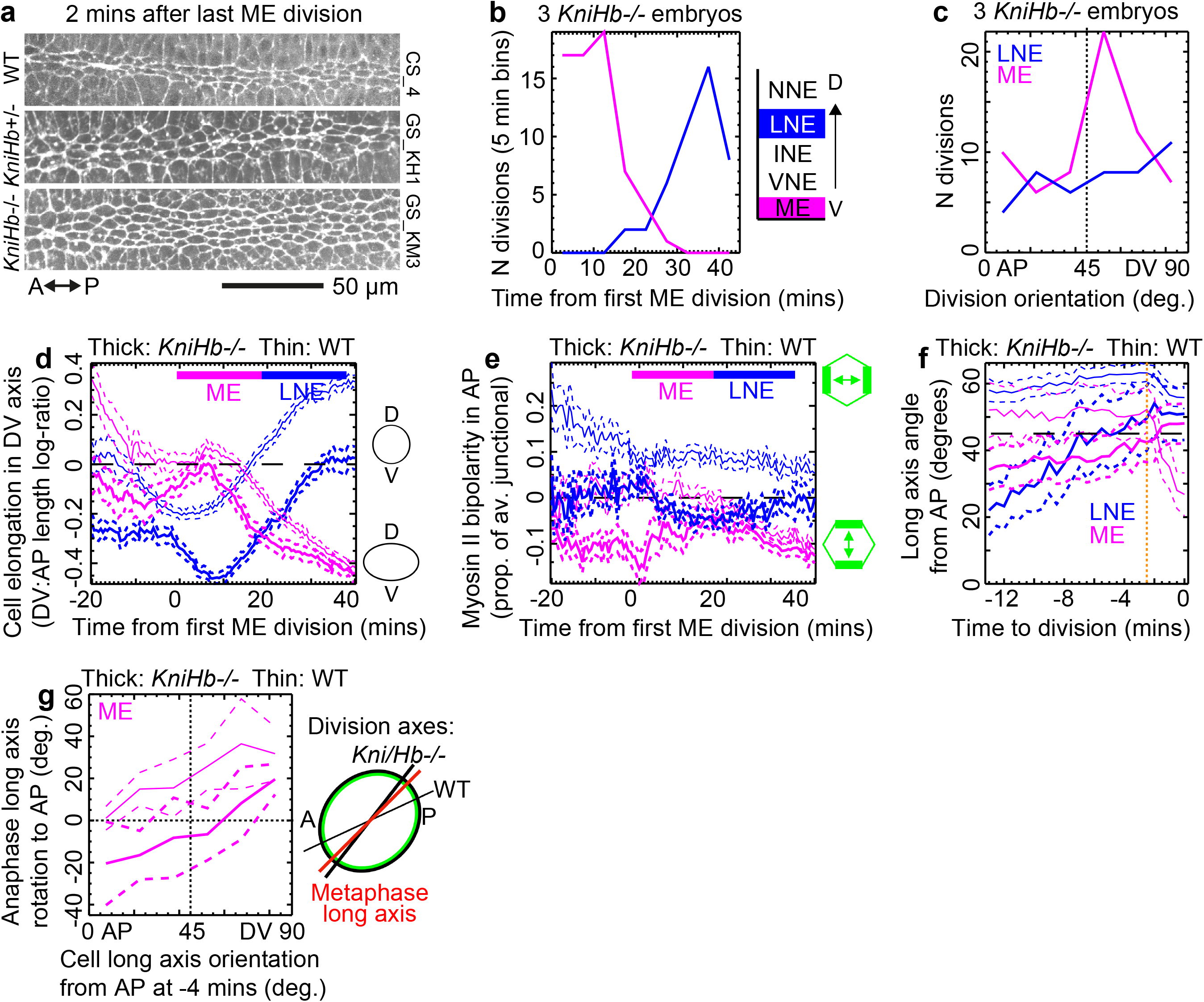
Anaphase long cell axis rotation towards AP is lost in an AP-patterning mutant. **a)** Stills from E-Cadherin channel for example embryos from three genotypes showing the width of the ME domain after division (see Fig. 6Sa). **b-g)** Data pooled from three *KniHb-/-* embryos. Comparable wild-type (WT) data for the same mitotic domains are presented as thin data lines. **b)** Frequency of divisions over developmental time. Bin size, 5 mins. **c)** Frequency of division orientation. Bin size, 15 degrees. **d-e)** Cell elongation (**d**) and average Myosin II bipolarity (**e**) over developmental time by mitotic domain. Coloured horizontal bars show domain division periods. **f)** Angle of cell long axis from AP during mitosis. Orange dotted line, start of anaphase. **g)** Rotation of cell long axis during anaphase versus metaphase (−4 mins) angle. Lines should intersect [x=45, y=0] if there is no rotation bias.

We tracked cells in *KniHb*-/-movies and aligned them in space and time as previously (Fig. 6Sb, c), identifying 65 ME and 45 LNE divisions across three embryos (Fig. 6b). Unlike WT, there was no AP-bias to the division orientation in ME (Fig. 6c). ME and LNE cells in *KniHb*-/-were more AP-elongated compared to WT (Fig. 6d and Fig. 6Sd), compatible with the posterior pull remaining intact whilst contractility in DV from polarised Myosin II is lost (Butler et al., 2009). Indeed, Myosin II bipolarity is mildly DV-oriented (negative) compared to AP-oriented (positive) in the WT (Fig. 6e). Importantly, in *KniHb*-/-embryos there is no long axis rotation to AP in anaphase in either ME or LNE divisions (Fig. 6f, g). Though not significant, there is a small rotation in the opposite direction, to DV in ME cells (Fig. 6g), compatible with the now AP-oriented (negative) Myosin II bipolarity through mitosis (Fig. 6Se).

Overall, when AP-patterning is eliminated, the long axis rotation to AP in anaphase is lost. We conclude that in WT, the anaphase rotation to AP is due to polarised Myosin II attracting the metaphase spindle.

### Compression from neighbouring dividing cells during metaphase re-orients divisions

Returning to the mechanical control of OCD, it remained a puzzle that the distribution of OCD in the NE (Fig. 4a) was not more bimodal, either AP-oriented due to AP-patterning (Fig. 5) or DV-oriented due to tissue stress (Fig. 4). Tissue stress anisotropy orients cells on average, but local stress can vary (Nestor-Bergmann et al., 2018). We observed that nests of quasi-synchronised cell divisions are common in cycle 14 *Drosophila* divisions (Fig. 7a and Supplementary Movies 1 and 6) (Foe, 1989), so we hypothesized that neighbouring divisions, as natural physical perturbations during mitosis, could re-orient OCD and explain the distribution of OCD in the NE. Just as the DV-oriented tissue stress in the NE may be temporarily reversed because of divisions in the domains on either side (Fig. 4j), tension across dividing cells could be reversed by neighbouring cells dividing on opposite sides (Fig. 7b) (Mao and Baum, 2015). We therefore asked whether neighbouring divisions re-orient each other.

**Figure 7:**
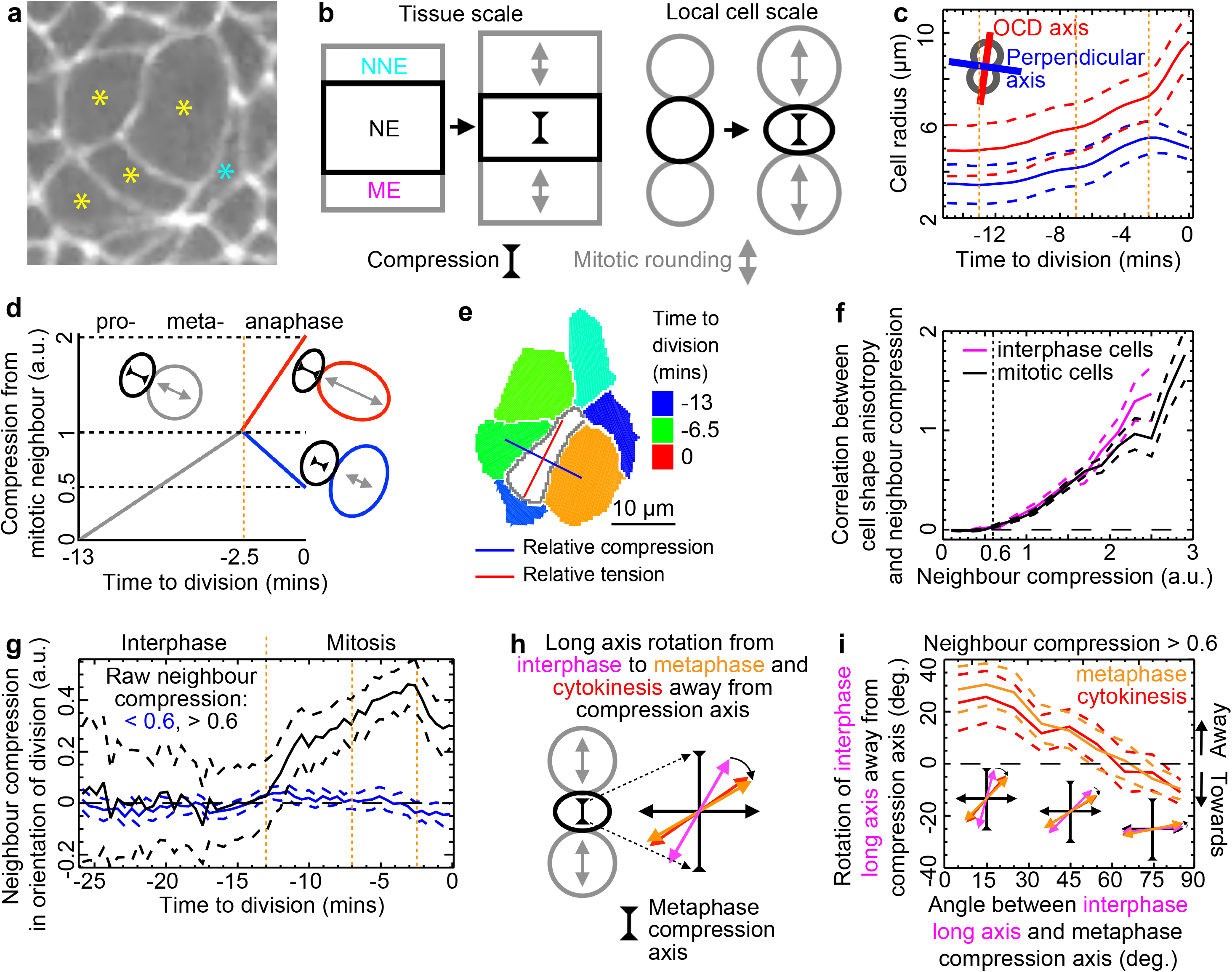
Compression from neighbouring dividing cells during metaphase re-orients divisions. **a)** Example nest of quasi-synchronised divisions (yellow stars) and their compressive effect on each other and neighbouring cells (blue star). **b)** Cartoon examples of anisotropic compression of neighbouring tissue (left) or neighbouring cell (right) divisions on focal tissue (NE, left) or focal cell (right). **c)** Cell radii in the future OCD (red) and orthogonal (blue) axes. Dashed lines are +/−SD. **d)** Schematic showing how time to division in a neighbouring cell (grey, red, blue) is converted to a compression on the focal cell (black), taking account of the relative orientation of the neighbouring division axis to the focal cell during anaphase. **e)** Example focal cell (white) surrounded by cells at various stages of mitosis. Large neighbours close to cytokinesis (orange and green) are on opposite sides, leading to a strongly anisotropic stress tensor (red and blue cross). **f)** Neighbour compression anisotropy starts to re-shape focal interphase and mitotic cells above a threshold value of 0.6. **g)** Strength of neighbour compression in the future OCD from interphase through mitosis. Positive values mean compression predicts perpendicular OCD. **h)** Scheme for how neighbour compression in metaphase should induce a rotation of the interphase cell long axis (magenta) away from the compression axis (here vertical) in metaphase (orange), thereby re-orienting the OCD (red). **i)** Re-orientation of the interphase cell long axis by neighbour compression in metaphase sets the OCD. Inset axes show approximate mean long axis rotations at x-axis values of 15, 45 and 75 deg. In all panels data are pooled from 5 Cad/Myo and 4 Cad/MT embryos. Except in (c), dashed lines are +/−95% CIs. Orange dotted vertical lines mark the onset of mitosis (−13 mins), metaphase (−7 mins) and anaphase (−2.5 mins).

Rounding up mitotic cells will release apical tension in surrounding cells, likely to the point of exerting compression on them (Fig. 7a) (Gupta et al., 2021), but we do not have access to the sign of stress. For simplicity, we will call this effect a compression. We are interested in the anisotropy of compression around each cell and its effect on the anisotropy of cell shape, which we know constrains the OCD. We therefore constructed a compression shear tensor for each cell to capture the anisotropic component of the effect of apically enlarging mitotic neighbours (see Methods). We made the assumption that the average enlargement of mitotic cell radii (Fig. 7c) linearly predicts the relative compression exerted on neighbours (Fig. 7d). The anisotropic compression tensor then summarises the compression of all neighbours, depending on their orientation and scaled by the length of shared interface (example in Fig. 7e).

We first tested whether the strength of the neighbour compression tensors was related to the strength of focal cell elongation. Indeed, these were positively correlated above a threshold compression strength of 0.6 for both interphase and mitotic cells (Fig. 7f, Fig. 7Sa). Remarkably, above this threshold, neighbour compression predicts OCD only during mitosis, most strongly at the end of metaphase (Fig. 7g), suggesting that neighbour compression in metaphase can re-orient the OCD. Note the drop in correlation in anaphase (> −2.5 mins), likely due to the influence of AP-patterning (Fig. 5).

For more direct evidence that neighbour compression re-orients OCD, we asked whether the interphase (−13 mins) long axis is re-oriented by metaphase (−3 mins) neighbour compression (Fig. 7h). We find that the cell long axis rotates towards the neighbour compression axis, up to an average of 30 degrees, depending on the angular difference between interphase long axis and neighbour compression axis (Fig. 7i, orange line). Furthermore, the subsequent division axis is set by this rotation (red line).

We conclude that anisotropic compression from dividing neighbours forces a new cell long axis orientation and that if this occurs during metaphase, this also sets the division axis.

## DISCUSSION

### Mechanics re-orients OCD during metaphase

Our work shows that, in the *Drosophila* embryo, epithelial cells can undergo a re-orientation of the axis of highest stress during metaphase, which results in a re-orientation of their OCD. Metaphase stress re-orientation occurs both due to a switch in tissue stress orientation and compression from neighbouring divisions.

We find that the absolute length of a cell’s shortest axis determines whether the mitotic spindle can rotate freely in the plane of the epithelium. Cell elongation can sterically constrain mitotic spindle movements. In the germband, therefore, stress orients the OCD indirectly through metaphase cell shape, though we cannot rule out that direct stress sensing is also in play (Lisica et al., 2022; Minc et al., 2011).

The control of planar OCD differs between tissues (Ramkumar and Baum, 2016; Tarannum et al., 2022; Taubenberger et al., 2020). During cycle 14 divisions the *Drosophila* germband is an immature epithelium with adherens but not septate junctions and lacking cell-ECM adhesions (Tepass et al., 2001). Active forces during cell division are therefore readily expressed as changes in cell shape. Germband neurectoderm cells are columnar, increasing their apical width by 50% before anaphase to accommodate planar rotation of the mitotic spindle. Even after mitotic rounding, the average cell elongation remains strong in metaphase (1.4 : 1), close to the steric hindrance threshold. By contrast, divisions in epithelia with stronger adhesions (Lisica et al., 2022) or ECM (Bosveld et al., 2016), or less columnar (Woolner and Papalopulu, 2012), show less dramatic deformations throughout mitosis and no steric hindrance. These may need other mechanisms to detect stress anisotropies.

### Temporary reversal of tissue stress anisotropy

DV tissue stress exceeds AP stress for most of the period we studied, resulting in DV-elongated cells. However, we show that there is a 15-min period of stress reversal when cells become temporarily elongated in AP. We exploited this reversal to show that only cell elongation in metaphase, when the spindle orientation is set, then sterically constrains the OCD.

Because Myosin II is no longer planar polarised to DV-oriented junctions in *KniHb-/-* embryos, the balance of stress is tipped to AP throughout. Nevertheless, a change towards more AP-elongated cells still occurs at the same time as the WT (Fig. 6d). A loss of tissue intrinsic DV tension due to Myosin II is therefore an unlikely cause of the stress reversal. A temporary release of tissue extrinsic DV tension is an alternative possibility. An increase in compliance of the dorsal amnioserosa tissue could contribute (Pope and Harris, 2008), but this would not explain why the loss of DV stress is temporary. We therefore favour the hypothesis that mitotic divisions in the ME and NNE domains flanking the NE in DV are directly responsible for the stress reversal, either releasing DV tension or actively compressing apices in DV, but more work is needed to confirm the nature of this stress change.

### Control of OCD by AP-patterning

We show that AP-patterning across the whole tissue, through the planar polarisation of Myosin II, biases the metaphase spindle towards AP for all mitotic cells. This generalises our previous finding that strongly Myosin II-enriched PSBs attract and stabilise spindle ends (Scarpa et al., 2018) suggesting that Myosin II could be directly involved in attracting the spindle, possibly through increased cortex stiffness (Spiro et al., 2014). AP-patterning straddles the whole DV circumference of the embryo (Irvine and Wieschaus, 1994) so that even the ME domain has polarised Myosin II, driving AP-oriented division in these isotropic cells. This polarity, and hence the spindle bias to AP, is lost in the *KniHb-/-*, replaced by the suggestion of an opposite bias to DV. It is possible that removal of AP-patterning reveals a less strong polarity in DV due to DV-patterning or mechanics.

Our results show that AP-patterning encodes a Myosin II-based mechanism that biases oriented cell divisions to AP and contributes to extending the AP body axis, joining forces with cell rearrangements. In vertebrate models, both cell rearrangements and cell divisions contribute to extending the AP axis during axis extension, with the loss of PCP randomising both rearrangements and divisions in Xenopus, zebrafish and mouse models (Morin and Bellaiche, 2011). The mechanism for how PCP ensures AP-oriented divisions has not been reported, nor how mechanics interacts with PCP, in these vertebrate examples.

### Interaction between mechanics and patterning

We find that discrepancies between the orientations of local stress and AP-patterning are resolved in metaphase by the positioning of the mitotic spindle, which in turn sets the perpendicular location of the cytokinesis ring and hence the OCD (Dimitracopoulos et al., 2020). Strong stress anisotropy at the tissue or local level can over-ride patterning to dissipate stresses through steric hindrance of the spindle orientation. Conversely, the extent of cell long axis rotation in anaphase reveals how much the stress axis has been over-ridden by patterning. Indeed, mildly elongated cells or cells with large apices readily orient their divisions to AP. In the posterior-most germband domain, cell elongation is on average AP-oriented, in the same orientation as AP-patterning, doubly ensuring AP-oriented divisions (da Silva and Vincent, 2007). Overall, AP-patterning on its own would ensure AP-oriented divisions, contributing to axis extension, but greater DV tension along most of the body axis over-rides patterning, leading to predominantly DV-oriented divisions.

The Hertwig rule, which states that the interphase cell elongation sets the OCD (Hertwig, 1884), is violated in three ways in the *Drosophila* germband: i) during the tissue stress reversal, stress anisotropy changes between interphase and metaphase, re-orienting the OCD; ii) compression from neighbouring cell divisions during metaphase re-orients the OCD, and iii) the spindle is attracted to planar polarised Myosin II in metaphase, orienting divisions to AP and away from the cell long axis. Our data suggests that the mechanical and patterning control of OCD in the germband does not require a memory of interphase cues.

## Acknowledgements

We acknowledge FlyBase (http://flybase.org/) for gene information and Dan Bergstralh, Nick Lowe, Daniel St Johnston, Bruno Monier, Stefano De Renzis and Bloomington Drosophila Stock Center (https://bdsc.indiana.edu/) for *Drosophila* strains and reagents. Laser ablations and their analysis were conducted with the Cambridge Advanced Imaging Centre. We thank Alex Nestor-Bergmann, Jocelyn Etienne, Claire Lye and other members of Bénédicte Sanson’s research group for useful discussions, and Part II student Hatty Cooper for her work at the beginning of this project. This work was supported by Wellcome Trust Investigator Awards to B.S. (099234/Z/12/Z and 207553/Z/17/Z). E.S. was also supported by a University of Cambridge Herchel Smith Fund Postdoctoral Fellowship and currently by a Royal Society Dorothy Hodgkin Fellowship (DHF_R1_201118). LM was supported by an EPSRC RSE fellowship (EP/R025398/1).

## Author contributions

G.B.B. conceived, developed and performed data and image analyses.

E.S. conceived, designed and conducted experiments, which E.S. and L.M. analysed.

G.B.B. prepared the manuscript, all authors revised it.

B.S. provided supervision and secured funding.

---

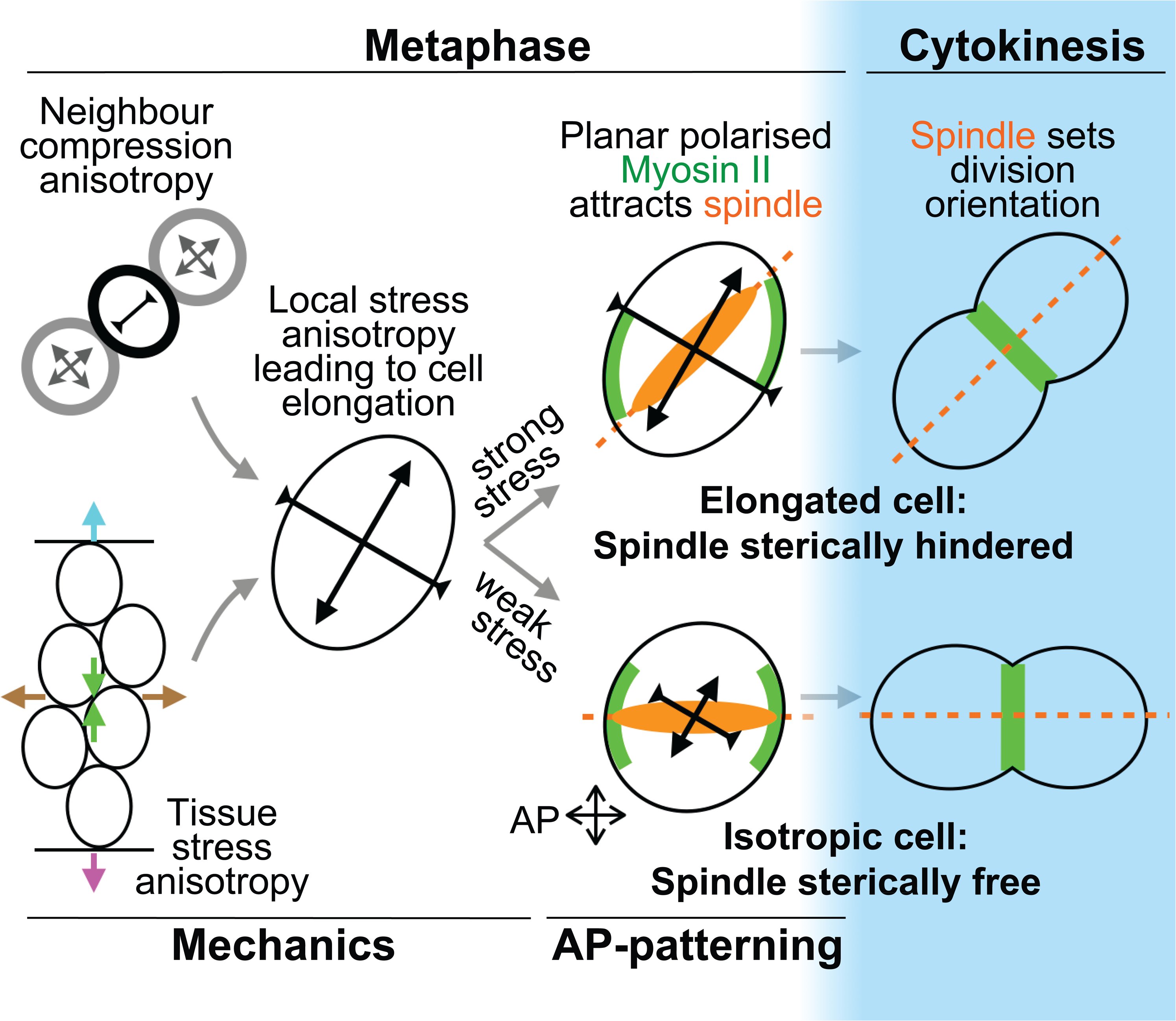

## METHODS

### 1. EMBRYO IMAGING

#### 1.1. Fly Stocks

Six *Drosophila melanogaster* stocks were used in this study. For wild-type live imaging, automatic tracking of cell divisions and fluorescence quantification we used *sqh^AX3^; shotgun::GFP* (Huang et al., 2009)*; sqh::mCherry* (Martin et al., 2009) that we refer to as ‘Cad/Myo’ and *ubi-E-Cadherin-GFP* (Oda and Tsukita, 2001)*; Jupiter::mCherry* (Karpova et al., 2006) that we term ‘Cad/MT’. We used the Anterior-Posterior(AP)-patterning knock-out *kni[10]hb[4*] (Butler et al., 2009)*; sqh:;GAP43mCherry* (Izquierdo et al., 2018)*.Sqh:EGFP[29B]* (Ambrosini et al., 2019) double mutant that we term ‘KniHb-/-‘ and heterozygote controls *t*hat we term ‘Kni/Hb+/−’.

*Sqh^AX3^; sqh-sqhGFP42;GAP43^mem^::mCherry/TM6B* (Tetley et al., 2016) was used for tissue-scale laser ablation and recombinant *Sqh-eGFP[29B]* (Ambrosini et al., 2019)*. Sqh::GAP43mCherry* (Izquierdo et al., 2018) was used for cut-out laser ablation experiments.

#### 1.2. Live imaging

Dechorionated embryos were transferred into halocarbon oil (Voltalef PCTFE, Arkema), mounted on a stretched oxygen-permeable membrane with their ventral side facing up and covered by a coverslip which was supported by a single coverslip bridge on either side of the membrane. Imaging was performed using a Nikon Eclipse E1000 equipped with a spinning disk unit (Yokogawa CSU10), laser module with 491nm and 561nm excitation (Spectral Applied Research LMM2), and a C9100-13 EM-CCD camera (Hamamatsu). Image acquisition was carried out using the Volocity software (Perkin Elmer). Imaging temperature was 21+/−1°C.

For live imaging, movie volumes ∼20 µm deep were taken for 50-100 mins covering cycle 14 divisions, always starting before the first mesectoderm (ME) division. Image volumes were sampled every 1 µm in depth and acquired every 30 s for Cad/Myo, KniHb+/- and KniHb-/-movies. Cad/MT image volumes were sampled every 0.7 µm in depth and every 20 s in time to image mitotic spindles and centrosomes. XY pixel size throughout is 0.364 µm with the AP field of view 254 µm from just posterior of the cephalic furrow (never in view, Fig. 1a), covering approximately three thoracic and the first two to three abdominal parasegments. Five wild-type Cad/Myo movies SC_1 to SC_5 are ‘080116’, ‘130116’, ‘160518’, ‘170518’ and ‘251116’, respectively, as used in (Scarpa et al., 2018).

### 2. TRACKING CELL SHAPES IN 2D AND 3D

#### 2.1. Cell tracking at and below the level of adherens junctions

Cell apices in movies were segmented and linked in time in an iterative process using an adaptive watershed algorithm (Blanchard et al., 2009; Butler et al., 2009) written in IDL (NV5 Geospatial). Briefly, the surface of the embryo was quantified as a smoothed ‘blanket’ spread over the apical-most junctional fluorescence in E-Cadherin movie channels. Cell segmentation and tracking was performed on curved planes at a constant depth relative to this embryo surface, with median and rolling ball filters applied as necessary to optimise subsequent cell tracking. For tracking at the level of adherens junctions (Ajs), a projection of local depths centred on 2-3 µm below the embryo surface was used (Fig. 1b, Fig. 1Sa and Table 1). For one Cad/Myo movie (CS_2) we also tracked cells below the level of Ajs at a depth of 7 +/−1 µm from the embryo surface (Fig. 4S2c). Manual corrections were further performed on movies CS_2 (at AJ and sub-AJ levels), CS_3, CS_4 and all KniHb-/-movies. Manual supervision focused particularly on dividing cells and their immediate neighbours.

**Table 1.** Algorithmic rules used to identify cell division events in cell tracking data.

| Rule | Description | Time window relative to last mother frame | Minimal gradient | Half-range |
| --- | --- | --- | --- | --- |
| 1 | Positive gradient of cell radius | 10 mins prior | 0.2 | 0.1 |
| 2 | Absolute cell radius | 0 mins | 7 | 3 |
| 3 | Positive gradient of the log-ratio of cell perimeter to cell area | 2.5 mins prior | 0.015 | 0.015 |
| 4 | Positive gradient of cell elongation log-ratio (long : short cell axes) | 2.5 mins prior | 0.1 | 0.1 |
| 5 | Positive gradient of cell dumbbell shape (see Fig. 1Sf, g) | 2 mins prior | 0.05 | 0.05 |
| 6A for Myosin II movies (CS_* and *KniHb*-/-) | Positive gradient of junctional Myosin II fluorescence density | 2.5 mins prior | 5 | 5 |
| 6B for MT movies (CM_*) | Negative gradient of whole cell MT | 3 mins prior | -2 | 1 |
| 7 | Negative gradient of cell radius | Between final mother frame and first daughter frame | -3.5 | 1.25 |

Cell segmentation and tracking was imperfect if Cadherin-attached fluorescence was locally weak; if the smooth epithelial surface ‘blanket’ was unable to follow cell apices into furrows, such as at the ventral midline before it was fully closed; and at the edges of images. We therefore removed partial cells at the movie edges and those with extreme or unlikely cell centroid jumps or cell shape changes (e. g. as seen in the first few frames of Supplementary Movie 1) from all subsequent analyses.

The number of cells successfully tracked over time per embryo is shown in Fig. 1Sd, e. For each cell at each time point, coordinates of cell centroids, perimeter shapes, bi-cellular (BCJ) and tri-cellular (TCJ) junction indices, and links forwards and backwards in time were stored.

#### 2.2. 2D cell shape analyses

Cell shapes were approximated by best-fit ellipses. Cell apices were first un-tilted and un-curved according to the surface normal and principal curvatures of the local tissue surface respectively. Best-fit ellipses were then found by minimising the area of mismatch between pixelated contours and ellipses, with constraints that the fitted ellipses have the same area and centroid as the cell contours.

AP and DV cell lengths (Fig. 4e, 4I, 4Sf, Fig. 6d, 6Sd) or cell lengths along the future orientation of cell division (OCD) (Fig. 7c) were calculated by projecting cell shape ellipses onto the relevant axis orientations, giving ellipse diameters in those orientations.

#### 2.3. Calculating 3D cell tilt and cell wedging between AJ and sub-AJ layers

To explore 3D cell shapes during the temporary tissue stress reversal, we calculated the tilt and 3D geometry of cells in the Cad/Myo movie CS_2. Having tracked cells at AJ and sub-AJ levels, we first needed to correctly match tracked cells between the two depths. We manually seeded five cell matches between the depths at the start and end of the movie. All remaining unmatched AJ cells were then sequentially matched to a sub-AJ cell using information from matched neighbours. The location of an unmatched sub-AJ centroid was predicted from the average AJ to sub-AJ vector of all neighbouring matched cells. The nearest actual sub-AJ centroid to this predicted sub-AJ centroid location was chosen as the match, providing it was within a threshold distance that we established to be half an apical cell radius. Progressively matching cells out from known matched cells filled out the whole tissue. We visually checked depth matches in a movie of the overlay of AJ and sub-AJ cell shapes with apico-basal centroid connections drawn (example in Fig. 4S2d).

Having matched cells between AJ and sub-AJ levels we were able to quantify 3D cell shapes, making the assumption that cell shape varies linearly between levels. For single cells, we measured 3D cell shape wedging and the tilt of the AJ centroid to sub-AJ centroid ‘in-line’ relative to the local surface normal (Fig. 4S2c). We calculated wedging and tilt quantities borrowing ideas from calculating morphogenetic strain rates (Blanchard et al., 2009). Instead of calculating rates of cell shape change and translation over time (per min), here we calculated these quantities over depth (per z µm) (Fig. 4S2c, e) (Deacon, 2012;

Sanchez-Corrales et al., 2018). The unit of wedging is proportional size change per mm in z, and tilt is a vector gradient (xy mm/z mm). We calculated rates of wedging in two orthogonal orientations, the principal axis being that in which there was greatest wedging, the minor axis perpendicular to this. Once calculated, cell wedging and tilt were projected onto our AP and DV embryonic axes. We used the convention that tilt and wedging change is from basal to apical, so an upright bottle-shape has negative cell wedging.

### 3. IDENTIFYING CELL DIVISION EVENTS

#### 3.1. Automated identification of cell divisions

We defined the completion of cytokinesis as when the mother cell splits into two daughter cells, marked by the appearance of a new segmented E-Cadherin interface between daughter cells. However, this alone is not a robust enough rule to identify cytokinesis events reliably because over-segmented cells can appear as two cells divided by a false interface. Through trial and error we evolved a set of algorithmic rules to identify cytokinesis events robustly in segmented and tracked cell data (Figure 1).

Each rule was computed for each cell in each movie frame and assigned a value between 0 and 1. These rules set the minimal change (gradient) in shape and fluorescence expected in the mother cell in the 10 mins prior to the completion of cytokinesis (see Fig. 1f). Gradients above the minimal gradient all evaluated to 1. For some rules this minimal gradient was effectively a threshold, with the rule evaluated to 1 above and 0 below. For other rules that were more likely to be broken, though in rare circumstances, the rule gradually evaluated to 0 with decreasing gradient according to a ‘half-range’ parameter, the gradient value at which the rule evaluated to 0.5 (Table 1).

A composite cell division probability is then calculated by multiplying all Rule values together. This method ensures that most but not all rules must be met, allowing for some unexpected division behaviour. Looking at distributions of the above Rule values for example divisions, along with visually minimising false positives and false negatives, we set a threshold composite probability of 0.01 above which cells were identified as dividing. We reasoned that missing data from the occasional missed division was less bad than including false data from the occasional false division.

To assess the success of this method we 1) visually confirmed that there were no false positives (see all cell divisions identified for embryo CS_4 in Supplementary Movie 2) and 2) compared cell divisions identified with the above algorithmic classifier with manually identified cell divisions.

#### 3.2. Manual identification of cell divisions

All visible cell divisions were manually located at the frame closest to cytokinesis completion in two movies in which cells had been tracked with automated methods only (CS_1 and CS_5) and two movies in which cell tracking had also been manually curated (CS_2 and CS_4). [x, y, frame] coordinates of manually identified cytokinesis events were compared to cell divisions identified with automated methods and the results shown in Table 2. Around 50% of divisions were correctly identified in the first two movies, rising to around 90% for manually curated movies. Failure to identify cell divisions was almost always due to interruptions in the tracking of mother cells in the prior 10 mins, or due to mis-tracked daughter cells immediately after division. In only a handful of cases did the above algorithmic rules fail to identify division when cells were correctly tracked. In the four movies we checked, with a total of 455 divisions, only 2 false positives were found.

**Table 2.** Comparison of manual to automated cell division classifications. Cytokinesis events were manually searched for in four Cad/Myo movies within the frame ranges indicated. No division events were identified in the first 20 frames of the movie since 10 mins (at 0.5 min frame interval) of mother cell behaviour was required to assess Rule 1 (see above). False negatives are divisions found manually but not identified with automated methods. False positives are events falsely classified as cell division by automated methods.

| Embryo Movie | Cell tracking method | N correctly identified | N false negatives | N false positives |
| --- | --- | --- | --- | --- |
| CS_1 (fr. 48-153) | Automated | 50 (44%) | 64 (56%) | 2 (2%) |
| CS_5 (fr. 102-229) | Automated | 54 (60%) | 36 (40%) | 0 (0%) |
| CS_2 (fr. 20-108) | Automated & manual curation | 60 (87%) | 9 (13%) | 0 (0%) |
| CS_4 (fr. 24-185) | Automated & manual curation | 171 (94%) | 11 (6%) | 0 (0%) |

The E-Cadherin channel in Cad/Myo movies was GFP attached to endogenous Shg whereas in Cad/MT movies it was GFP attached to over-expressed ubi-E-Cadherin. As a result, the E-Cadherin signal was stronger and automated cell tracking was more reliable in the Cad/MT movies, so manual curation was not necessary. GFP in *Kni/Hb-/-* movies was attached to endogenous Shg, as in Cad/Myo embryos, so were all manually curated.

#### 3.3. Cell division tracking summary

With the above methods we identified 450 divisions in five Cad/Myo movies, 286 divisions in four Cad/MT movies and 122 divisions in three *KniHb-/-* movies. At the sub-AJ level, we identified 67 mitosis events (Fig. 4S2a, b) in embryo CS_2 at the same times and covering the same NNE, LNE and INE division domains as for AJ level tracking (Fig. 2Sh). All cell divisions were cycle 14 divisions captured between 0 and 100 mins after the first ME division, straddling stages 8 to 10 (Bate and Martinez Arias, 1993).

Each dividing cell was then assigned its own ‘time to division’ axis, with the frame after which the mother splits into two daughters set to 0 mins.

### 4. EMBRYONIC TIME & SPACE AXES

Embryo movies were co-aligned according to a set of temporal and spatial developmental landmarks so that data from different movies and different genotypes could be overlaid and compared directly.

#### 4.1. Movie staging and synchronisation

The temporal developmental origin (0 mins) in all movies of all genotypes was set to the completion of the first observed mesectoderm (ME) cytokinesis. This is equivalent to approximately 35 mins after the start of germband extension (Butler et al., 2009; Tetley et al., 2016) and marks approximately the transition from fast to slow phase of germband extension (Irvine and Wieschaus, 1994) as the cell intercalation strain rate slows down (Butler et al., 2009).

We used all data between −20 (mid stage 7) and 100 mins (mid stage 10) in our WT movies, each of which covered a subset of this range (Fig. 1Sd, e). Within this range, approximately 0 to 30 mins represents stage 8 and 30 to 75 mins is stage 9 (Bate and Martinez Arias, 1993). Note that our imaging temperature of 21°C is lower, hence development slower, than most early staging reports that were imaged at 25°C. Because AP-patterning gene expression changes from primary pair-rule (stage 7) to secondary pair-rule (stage 8) to segment polarity genes (stages 9-10) over this time range (Clark and Akam, 2016), we were careful throughout all analyses to be clear from which stage data is presented.

#### 4.2. Anterior-posterior (AP) coordinates

We were interested in where cell divisions occurred within each parasegment because the neurectoderm has previously been split into mitotic domains with different within-parasegment AP locations (domains M, N, 21, 25 in the anterior trunk (Foe, 1989)). Parasegment boundaries (PSBs) were identified in Cad/Myo movies as the strong periodic DV-oriented cables of Sqh-GFP (Fig. 2Se). Groups of cells in between strong Sqh-GFP enrichments were manually selected (each group corresponding to a single parasegment) at various points during each movie. Because cells were tracked over time, these classifications of parasegmental group identity could be automatically back- and forward-tracked to define the same groups of cells throughout the movie. Cells were allocated to numbered parasegments starting with zero for the most anterior parasegment in view (Fig. 2Sf). Because the cephalic furrow was not in view, these parasegement numbers were approximate and we were not able to distinguish between even and odd parasegments. A within-parasegment coordinate was assigned to each cell, normalised to between 0 at the anterior PSB and 1 at the posterior PSB, as in (Tetley et al., 2016) (Fig. 2Sg and Supplementary Movie 3). We found that a threshold within-parasegment coordinate of 0.75 separated anterior neurectoderm divisions from a distinct posterior population that divided approximately between 40 and 60 mins (Fig. 2b, c).

#### 4.3. Dorso-ventral (DV) coordinates and domains

DV cell location was set in µm from the ventral mid-line (VML) at the end of DV tissue convergence (+15 mins, Fig. 2Sb). Cells carried these DV coordinates forwards and backwards in time, defining a co-moving coordinate system. This coordinate system was in turn used to classify five separate cell-type domains in DV in each movie so that we could pool cells from different movies into the same cell types for analysis.

The most ventral cells were easily distinguished as mesectoderm (ME) cells by their early cell divisions (complete by 20 mins) and small AP-oriented shapes thereafter. The ME occupied a region approximately 0-12 µm from the VML. On the dorsal side of the neurectoderm (NE), the non-neural ectoderm (NNE) cells were identified from their early cell divisions which were also complete by 20 mins. A line at 66 µm from the VML separated the NNE from the NE (Fig. 2Sc, d).

The NE was broken down into three domains, ventral (VNE), intermediate (INE) and lateral (LNE), that from cell counts of in situ images of *ind* expression (Lye, Blanchard, Nestor-Bergmann, Evans & Sanson, in prep.) map approximately to *vnd*, *ind* and *msh* expression domains. *vnd* cells are unique in becoming progressively elongated in DV and larger, occupying a band between the border with the ME at 12 µm and 30 µm. We distinguished *ind* from *msh* domains from their different cell division timings (mitotic domains 21 and N in (Foe, 1989), respectively). This allowed us to identify the approximate dividing line at 48 µm from the VML.

To measure angles relative to embryonic axes, the orientation of the ventral midline over time for each movie was used as the evolving orientation of the AP axis. Note that some embryos rolled a little in the DV axis in the field of view, but this is corrected for by keeping track of the ventral midline as it rolled.

#### 4.4. Landmarks of mitosis

In Figure 3e we relate our measured average changes in morphology and fluorescence during mitosis to known mitotic landmarks and checkpoints as reviewed in (Ramkumar and Baum, 2016). We do not know when the G2/M (interphase/prophase) transition occurs in our data, corresponding to the rise in CDK1-cyclin B activity, but our first measurable change associated with mitosis is a drop in apico-medial Myosin II at −13 mins. We therefore call this the effective onset of prophase in our data.

Prometaphase is defined as starting with nuclear envelope breakdown (NEBD) and ends with spindle formation as metaphase commences. We see two Jupiter-mCherry centrosomal foci below each nucleus during prophase, which disappear at around −7.5 mins, which we define as NEBD. In a separate study, NEBD was also seen to occur at approximately −7 mins in the germband from the loss of mTor-YFP, a nuclear envelope marker (Lye et al., 2014). A variably oriented elongated spindle then appears almost immediately, so we set the onset of metaphase in our data to −7 mins.

Passing through the spindle assembly checkpoint (SAC) is defined by the onset of spindle separation, with associated cell elongation, and marks the start of anaphase. We see that anaphase cell elongation starts at −2.5 mins and the cytokinesis ring appears at −2 mins in our data so we set the SAC to −2.5 mins.

### 5. LASER ABLATION

#### 5.1. Tissue scale and cut-out box ablations

Laser ablation experiments were performed using a TriM Scope II Upright 2-photon Scanning Fluorescence Microscope controlled by Imspector Pro software (LaVision Biotec) equipped with a tuneable near-infrared (NIR) laser source delivering 120 femtosecond pulses with a repetition rate of 80 MHz (Insight DeepSee, Spectra-Physics). The laser was set to 927nm, with power ranging between 1.05-1.70 W. The maximum laser power reaching the sample was set to 220 mW and an Electro-Optical Modulator (EOM) was used to allow microsecond switching between imaging and treatment laser powers. Laser light was focused by a 25x, 1.05 Numerical Aperture (NA) water immersion objective lens with a 2mm working distance (XLPLN25XWMP2, Olympus). Ablations were carried out during image acquisition (with a dwell time of 9.27 µs per pixel), with the laser power switching between treatment and imaging powers as the laser scanned across the sample. For tissue-scale cuts, targeted line ablations of 20 µm length were performed in an AP or DV orientation using a treatment power of 220 mW on the *vnd* domain (rows of cells 2-3-4 from the ventral midline) of the ventral neuroepithelium of stage 9 embryos. Embryos were staged by waiting for midline closure after the last division in the ME (mitotic domain 14), equivalent to 25-30 mins after the first ME cell division. Images were acquired with a frame delay of 1 s for tissue-scale cuts and 753 ms for cut-out ablations.

#### 5.2. Analysis of ablation recoil velocity

To quantify recoil velocity, we automatically detected the position of the ablation line in the images, chose a region of interest around it, detected the fluorescent structures inside this region and inferred their motion from the video sequences. The region of interest in each movie is defined as a 30 pixel area on each side of the cut. The position of the ablation line was recorded by taking a screen snapshot at pre-acquisition. The screenshot was cropped and the site of cut automatically matched to its location in the video sequence. To analyse recoil velocities, we first performed two pre-processing steps: a frequency filter to remove stripes caused by electrical interference in the detector and a median filter denoising step. Subsequently, we computed the optical flow in order to estimate the velocity of the structures using the algorithm described in (Brox et al., 2004) in the whole region of interest and on the bright pixels of signal in the region of interest for each embryo. In order to use the flow of Myosin II to measure recoil away from the cut, only the velocity component of flow perpendicular to the cut is considered in the analysis. For each side of the cut, the average normalised relaxation speed is computed and the two averages are combined to give a final normalised relaxation speed away from the cut (Fig. 4g). This final velocity was compared between the two conditions with a ‘Two sample t-test’.

#### 5.3. Analysis of ablation cell shapes

For cut-out laser ablation, cell shapes were analysed 5 timepoints (−3.9 s) before ablation and 20 timepoints (15.06 s) after ablation. The outline of each cell was manually traced over the Sqh-eGFP channel using the polygon tool in Fiji and its best fit ellipse was calculated. Shape descriptors (long axis, short axis, log-ratio of long:short axis) were calculated from the best fit ellipse while the area of cells were measured from manually traced polygons. The AP axis of each embryo was located by taking a low magnification image.

### 6. FLUORESCENCE INTENSITY

#### 6.1. Fluorescence normalisation

For each cell in each movie frame we quantified the density of fluorescence in each movie channel associated with membrane (junctional) and cytoplasmic (medial) domains. Bleaching of Sqh-mCherry, Shg-GFP, E-Cad-GFP and to a lesser extent Jupiter-mCherry, was significant over the 50-100 mins of each movie, so we normalised the greyscale range of each channel in each movie frame. For example, for Shg-GFP, we set the lowest 20% of pixels as background (0 greyscale) and pinned the 99.5^th^ percentile to 200 greyscale, allowing the top 0.5% to occupy the top 55 greyscale with minimal overexposure at 255. Background and signal percentiles for all movies are shown in Table 3.

**Table 3.**
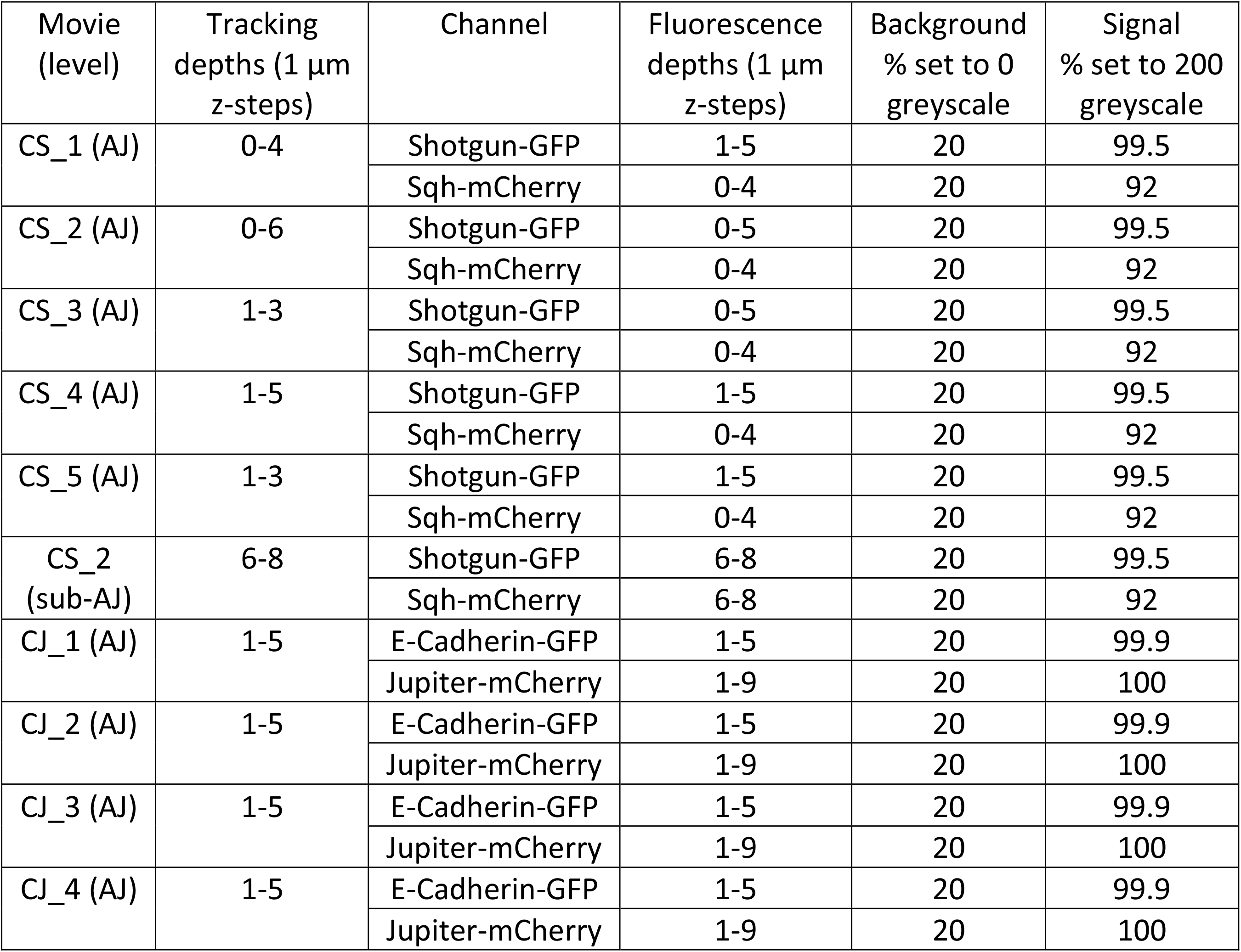
Wild-type embryo movie details. Projection depth ranges for tracking and fluorescence quantification are relative to the curved embryo surface. Fluorescence in each movie volume image is normalised to 0-200 greyscale between ‘background’ and ‘signal’ percentiles.

#### 6.2. Junctional and medial fluorescence quantification

Junctional and medial domains were determined by the outline coordinates of each cell, as recorded by cell shape tracking. The coordinates of non-overlapping pixelated cell perimeters were stored by the cell tracking for each cell in each movie frame. The fluorescence intensity of a particular movie channel at cell perimeters was quantified as the average of cell perimeter pixels and neighbouring pixels to a width *w* = 0.65 µm perpendicular to the local perimeter, meaning the perimeter pixel and one pixel either side. For the Myosin II channel, this was a compromise between being wide enough to encompass all junctional Myosin II fluorescence but narrow enough to minimise the inclusion of medial Myosin II. We then summarised the fluorescence intensity of every cell-cell junction as the average intensity of all perimeter pixels along the junction.

Fluorescence density at TCJs is expected to be 1.5 times that of BCJs because of the convolution of the confocal point spread function of three junctions meeting. This extra fluorescence at TCJs contributed by the neighbouring cell BCJ pollutes attempts to summarise the polarity of membrane fluorescence, particularly in elongated cells in which vertices tend to be clustered towards the cell long ends (Bosveld et al., 2016; Scarpa et al., 2018). We defined TCJs as pixels within 0.65 µm radius of the TCJ centre point, allowing us to remove these and define a ‘no-vertex’ BCJ density (see ‘Top-down view’ in Fig 5Sg). For Myosin II channels, the fluorescence density of each cell-cell interface in each movie frame was calculated as the average intensity of ‘no-vertex’ BCJ pixels.

#### 6.3. Portraits of average cell fluorescence through mitosis

We set out to build up a portrait of the behaviour (shape and fluorescence) of the average cell in 3D through mitosis. To do this, we needed to overlay and orient cells in the same way so that we could average the fluorescence surrounding cells in both movie channels in Cad/Myo and Cad/MT movies.

We first super-imposed all cells by translating each cell’s centroid (from AJ level tracking) to the origin, bringing along both the local embryo surface normal and a local 3D volume of fluorescence from both movie channels. Secondly, we rotated the local fluorescence volumes so that each cell’s local surface normal aligned with the positive z-axis, meaning that cell bases extended from the cell centroid in negative z. Finally, we rotated the volumes around the z-axis to set the future OCD to the x-axis. With all cells overlaid, standing upright and dividing along x, we were able to take the average voxel fluorescence intensity in the surrounding image volumes. 411 ectoderm cell divisions were overlaid from Cad/Myo embryos, with 270 from Cad/MT embryos.

Cutting this averaged fluorescence volume at z=0 gives the shape and fluorescence of the averaged cell at the AJ level (top row of Fig. 3c, d and 2^nd^ panel of Supplementary Movies 4 and 5). Cutting at z=4 shows the average cell behaviour at a sub-AJ level (bottom row of Fig. 3c, d and 4^th^ panel of Supplementary Movies 4 and 5). Individual mitotic cell behaviour is variable (see Fig. 3Se and Supplementary Movie 2). This method yields smooth average mitotic behaviour, separated into clear medial and junctional fluorescence, with horizontal elongation in the future OCD and a vertical cytokinesis ring. The similarity in Shotgun and E-Cadherin fluorescence patterns between the Cad/Myo and Cad/MT average cells allows comparison of the Myosin II with the MT patterns at different depths.

#### 6.4. Myosin bi- and unipolarity

To quantify the apical junctional planar polarity of Myosin II, we used the ‘no-vertex’ BCJ measures from above. We first expressed BCJ fluorescence intensity around each cell perimeter as a function of planar angle around the cell centroid. Treating this intensity signal from 0 – 360 degrees as a periodic repeating signal, we calculated its Fourier decomposition, extracting the amplitude of the period 2 component as the strength of Myosin II bipolarity (equivalent to planar cell polarity), with its phase representing the orientation of cell bipolarity (Fig. 6h, 6Sh). We also extracted the period 1 component as a Myosin unipolarity measure (Fig. 6Si). We normalised the strength of both polarities by dividing the Fourier amplitudes by the mean cell perimeter Myosin II signal, so that they would be consistent across images, frames and embryos.

Tensors describing the orientation and strengths of the principal axes of Myosin II bipolarity were projected onto embryonic axes in a similar manner to the ‘2D cell shape analyses’ section above.

Using the above methods we produced uni- and bipolarity measures projected along the AP axis for each cell at each time point, that were independent of each other and normalised to control for variation in Myosin II fluorescence.

#### 6.5. Jupiter fluorescence quantification

We quantified average Jupiter fluorescence at junctions and medially across depths 1–9 µm from the tissue surface (Fig. 6d, e).

#### 6.6. 3D spindle tracking

To quantify spindle depth and rotation accurately, we applied an adaptive binary fluorescence threshold to distinguish Jupiter pixels from Jupiter-free cytoplasmic pixels (Fig. 3Sc, Fig. 5Sa). Principal component analysis (PCA) of the 3D clouds of thresholded Jupiter pixels provided ellipsoid fits to spindle shapes for each cell (Fig. 5c). The shape of the spindle identified in this way was essentially linear, dominated by its principal component, with much smaller widths orthogonal to the spindle long axis (Fig. 5Sb).

### 7. Constructing an anisotropic neighbour compression stress tensor

Cell rounding in prophase and metaphase, and cell elongation during anaphase, are actively driven by increased cortical actomyosin activity in dividing cell (Ramkumar and Baum, 2016; Rosa et al., 2015). The rounding and elongation would therefore be expected to exert outward forces on neighbouring cells, compressing them. Cells under isotropic planar compression can be forced to divide out of the plane but we were more interested in anisotropic compression that would influence planar cell elongation in metaphase, constraining the spindle orientation and hence the OCD (Fig. 5).

We therefore set out to construct a summary measure of the anisotropic compression that cells are under from surrounding mitotic cells. We did not have direct access to differential cell pressures in our live movies so instead made the simplifying assumption that the enlargement of cells during mitosis is a proxy for the strength of compression they exert on neighbours. That is, we assumed that the active rounding force increased with how rounded cells became through prophase to metaphase. During anaphase, elongation along the future division axis also exerts a pressure in the direction of elongation (Gupta et al., 2021), less so perpendicularly as the cytokinesis ring actively contracts the dividing cell. We used the approximately linear increase in radius of cells in prophase to metaphase and the elongation of cells in the OCD during anaphase (Fig. 7c) to estimate the likely compressive force exerted (Fig. 7d).

Then for each cell in interphase and mitosis we constructed a compression tensor by adding the compressive effect of all neighbours, in the orientation of each neighbour and scaled by the length of the shared interface. To extract the anisotropic part of the compression tensor we subtracted the isotropic part or Trace. The resulting anisotropic tensor was negative in the principal axis of relative compression, positive in the orthogonal axis of relative tension (see Fig. 7e).

---

**Figure 1 Supplement.**
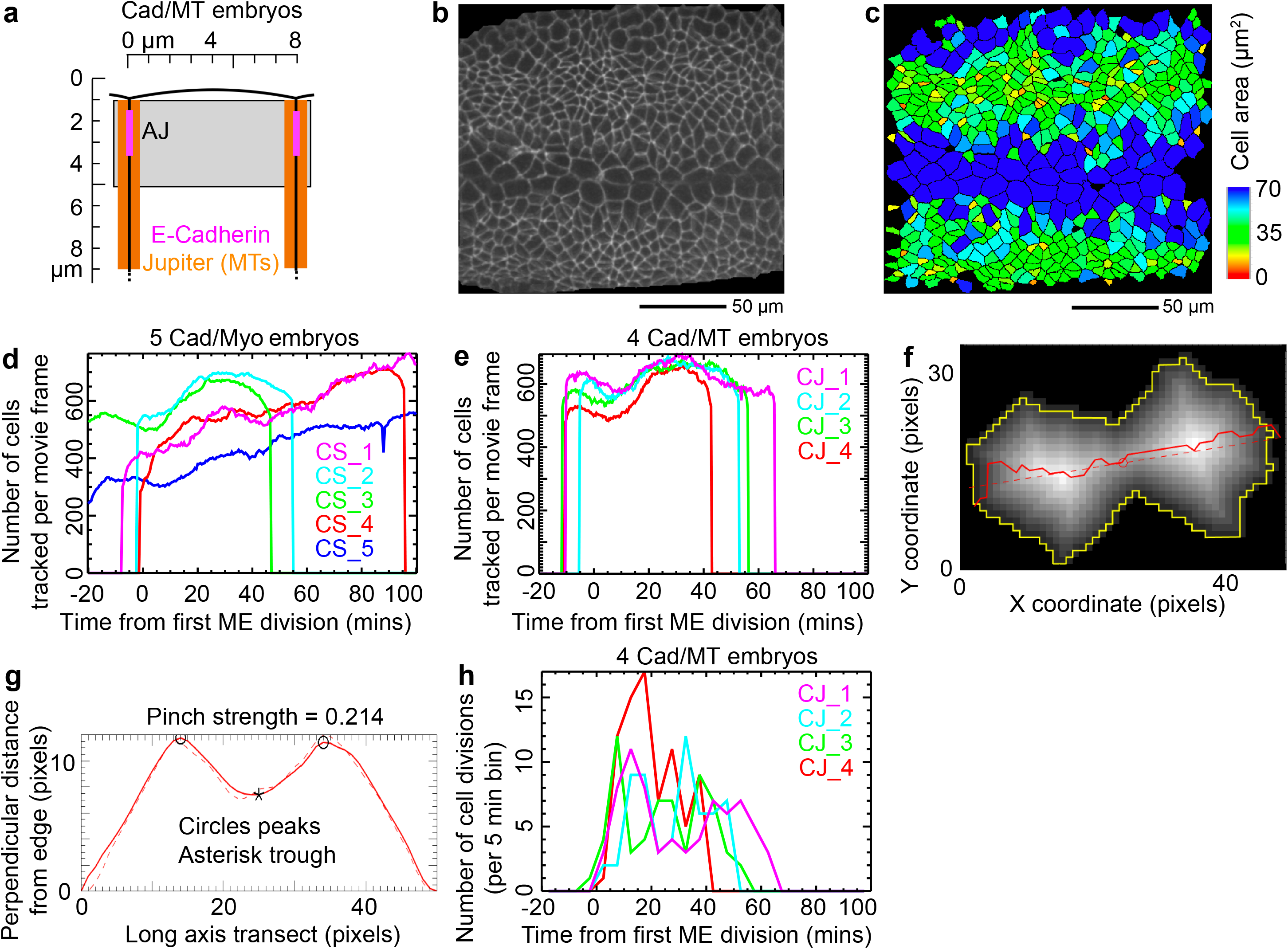
Tracking cell divisions *in vivo*. **a)** Schematic of epithelial cell apex showing the depth projection range (grey box) relative to cell apices used to track cells at the level of AJs in Cad/MT movies. **b)** Projection of E-Cadherin channel at the level of AJs at 0 mins from Cad/MT movie CJ_3. Central horizontal band of large cells are mesectoderm (ME), straddling the ventral midline (VML). The rest are ectoderm cells, with non-neural ectoderm cells dividing at the top of the image. Anterior left. **c)** Segmented cell shapes in (**b**), colour-coded by surface area. **d)** Number of cells segmented and tracked successfully per image frame per Cad/Myo movie. **e)** Same as (d) for Cad/MT movies. **f, g)** Calculating ‘dumbbell pinch’ measure for an example cell outline (yellow line). A mid-line path (red line) is calculated as the watershed of the distance to outline map (greyscale) along the long axis of the cell (**f**). This path is simplified to the regression line fit to its path points (dashed red line). The distance from pixel outline along this regression path has two peaks for a dumbbell shape (**g**). The difference in height between the lesser of the two peaks and the minimum between peaks, expressed as a proportion of the former, is used to quantify the ‘dumbbell pinch’ measure. **i)** Frequency of cell division events over developmental time for four Cad/MT embryo movies.

**Figure 2 Supplement.**
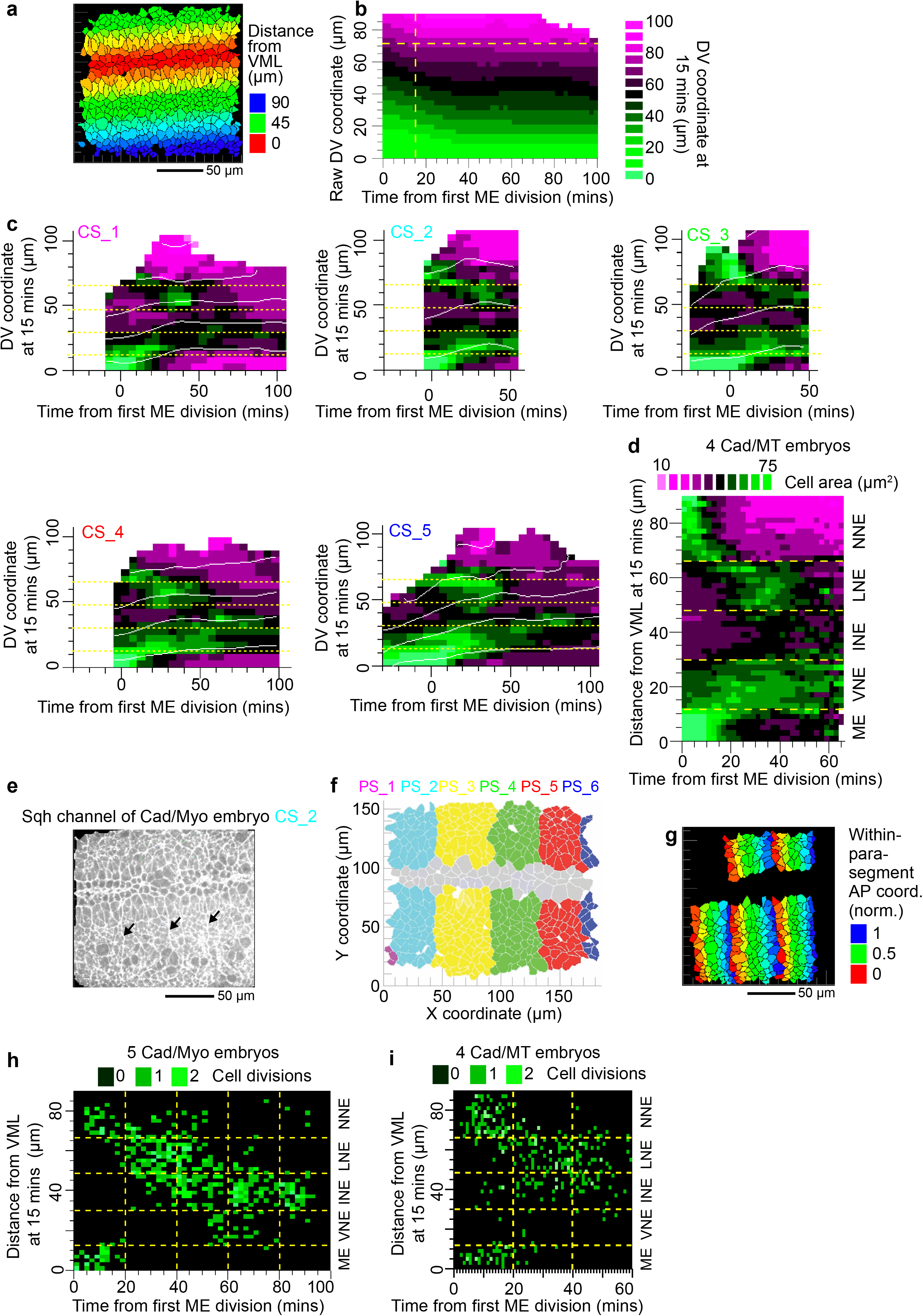
Assigning divisions to known mitotic domains. **a)** DV co-moving coordinates assigned to cells at 15 mins (embryo CS_2). Red (DV=0) shows ME cells at the VML. **b)** Flow of cells in DV across time showing that convergence stops in lateral tissue at 15 mins after the first VML division (dashed yellow line). Data pooled from 5 Cad/Myo embryos. **c)** Fig. 2a broken down by embryo. **d)** Average cell areas for pooled data from four Cad/MT embryos. Yellow dashed lines show manually assigned borders between DV domains with different patterns of cell shape, as in Fig. 2a. Y-axis is a DV co-moving coordinate frame, assigned at 15 mins then constant for each cell over time. **e)** Fluorescently tagged Myosin II (sqh) at AJs at 15 mins (embryo CS_2). Arrows highlight parasegmental boundaries (PSBs). Anterior left. **f)** Example movie frame showing manually assigned cell membership of parasegments and ME cells (grey). **g)** Cells colour-coded by normalised within-parasegment AP coordinates, assigned between neighbouring PSBs (same image frame as in (**e**)). Cell dataset is reduced because boundaries on both sides of a parasegment are required to assign a within-parasegment coordinate. **h)** Spatio-temporal map of identified division events pooled from five Cad/Myo embryos. Horizontal yellow lines show co-moving DV boundaries (as in Fig. 2a) used to classify cells into mitotic domains. Vertical lines show 20-min phases separated in Fig. 2b. **i)** Same panel as in (**h**) but for four Cad/MT embryos.

**Figure 3 Supplement.**
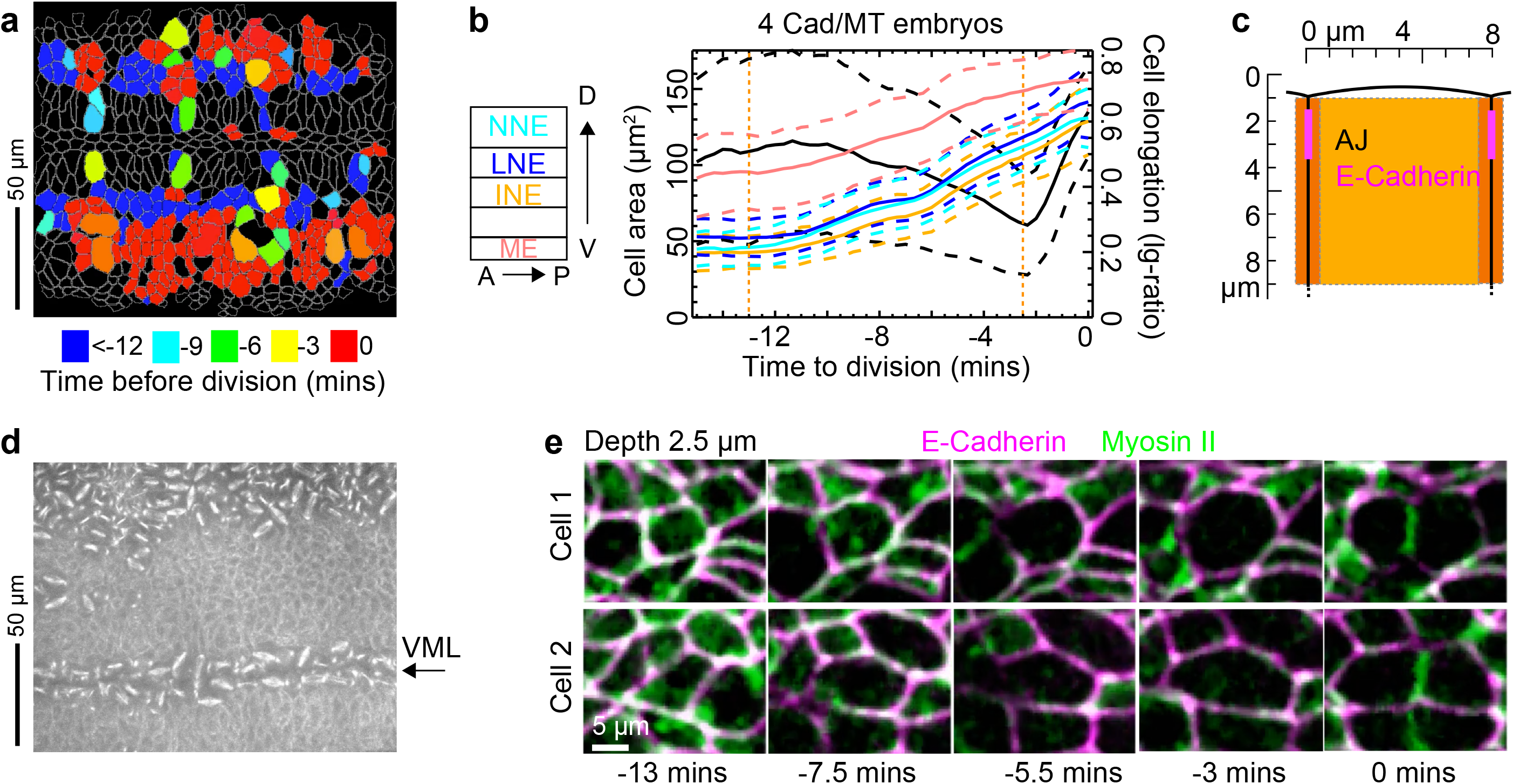
The time course of mitosis. **a)** Still from Supplementary Movie 1 (bottom right panel), showing cells classified as dividing during embryo movie CS_4, colour-coded by time before division at 49 mins. Cells in red have divided (NNE, ME, LNE and some WG), whilst blue cells (IND and some WG) will divide within the next 50 mins. Anterior is left. **b)** Cell shape through mitosis at the level of AJs for Cad/MT embryos. Left y-axis, cell area for four mitotic domains. Right y-axis, cell elongation ratio (black line). **c)** Schematic of epithelial cell profile showing the depth projection range (light orange box) relative to cell apices used to quantify cytoplasmic MT density in Cad/MT movies. **d)** Projection of the depth range in (c) from MT channel of Cad/MT movie CJ_4 at 10 mins. Lower horizontal band of mitotic spindles are ME cells straddling the VML (arrow), top band is NNE. LNE cells in between are starting to enter mitosis. Anterior right. **e)** Average cell fluorescence around the cell centroid for two example Cad/Myo cells at the level of AJs, 2.5 µm from the surface of the epithelium. Averaged data pooled from hundreds of cells are shown in Fig. 3c, d and Supplementary Movies 4 and 5.

**Figure 4 Supplement 1.**
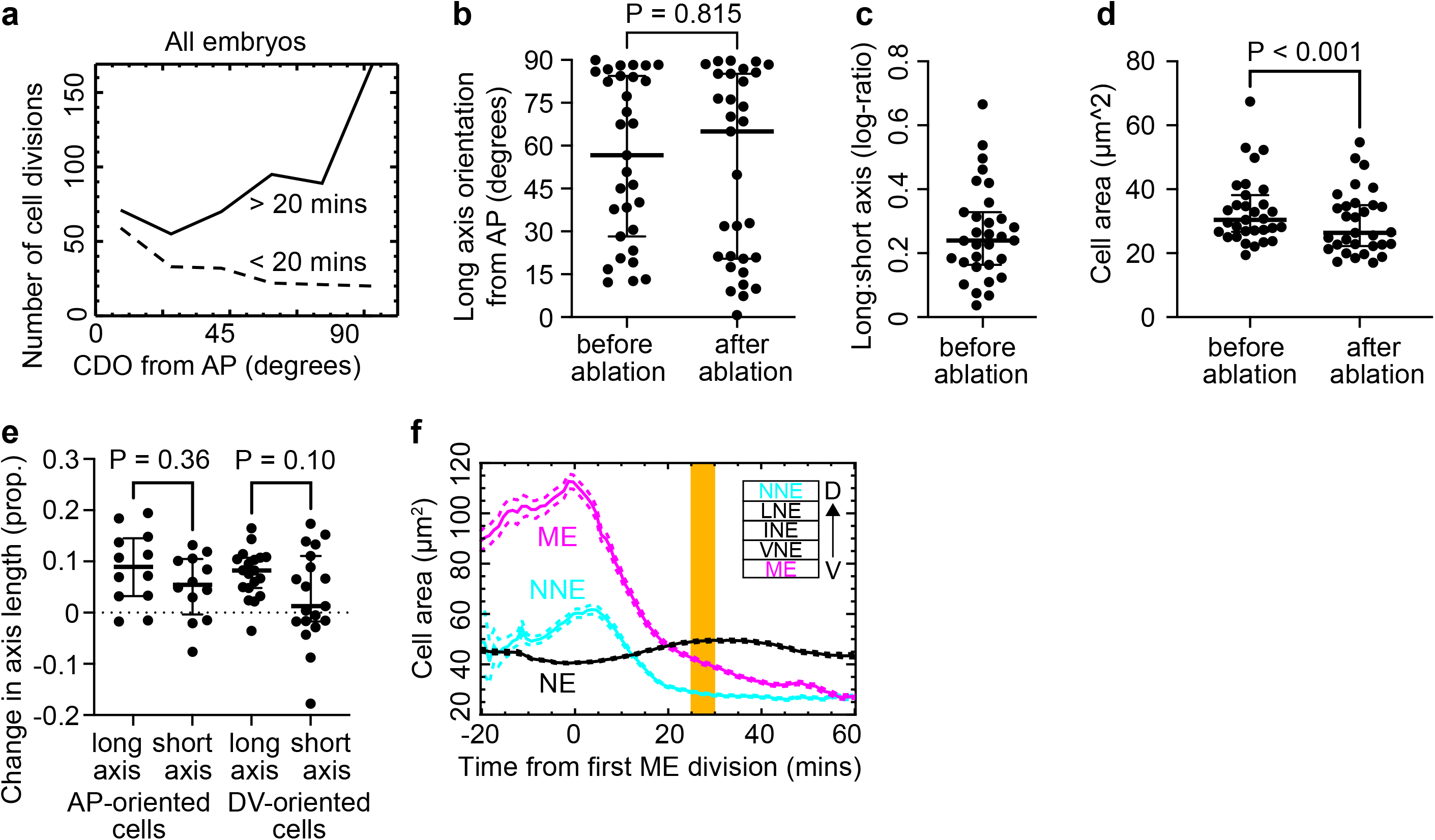
Metaphase cell shape tracks tissue tension. **a)** OCD for early (<20mins, mostly ME and NNE) and late (>20mins, mostly NE) divisions. Bin size is 15 degrees. **b-e)** Box laser cut results. **e)** Distribution of the orientation of cell long axes before and 15 s after ablation, showing that a range of AP- and DV-oriented cells were selected, though it was difficult to find cells aligned perfectly with AP (Mann-Whitney test, N = 31). **f)** The range of eccentricities before ablation of cells selected for box ablation (N = 31). **g)** Reduction in cell areas after box cut ablations (Paired t-test; N = 31). **h)** Proportional reduction in axis lengths after box ablation, broken down by orientation of cell long axis before ablation (Paired t-tests; AP-oriented, N = 12; DV-oriented, N = 19). **i)** Apical cell areas in ME, NNE and pooled NE mitotic domains over developmental time (see Fig. 4e). Tissue laser cuts were performed at 25-30 mins (orange bar).

**Figure 4 Supplement 2.**
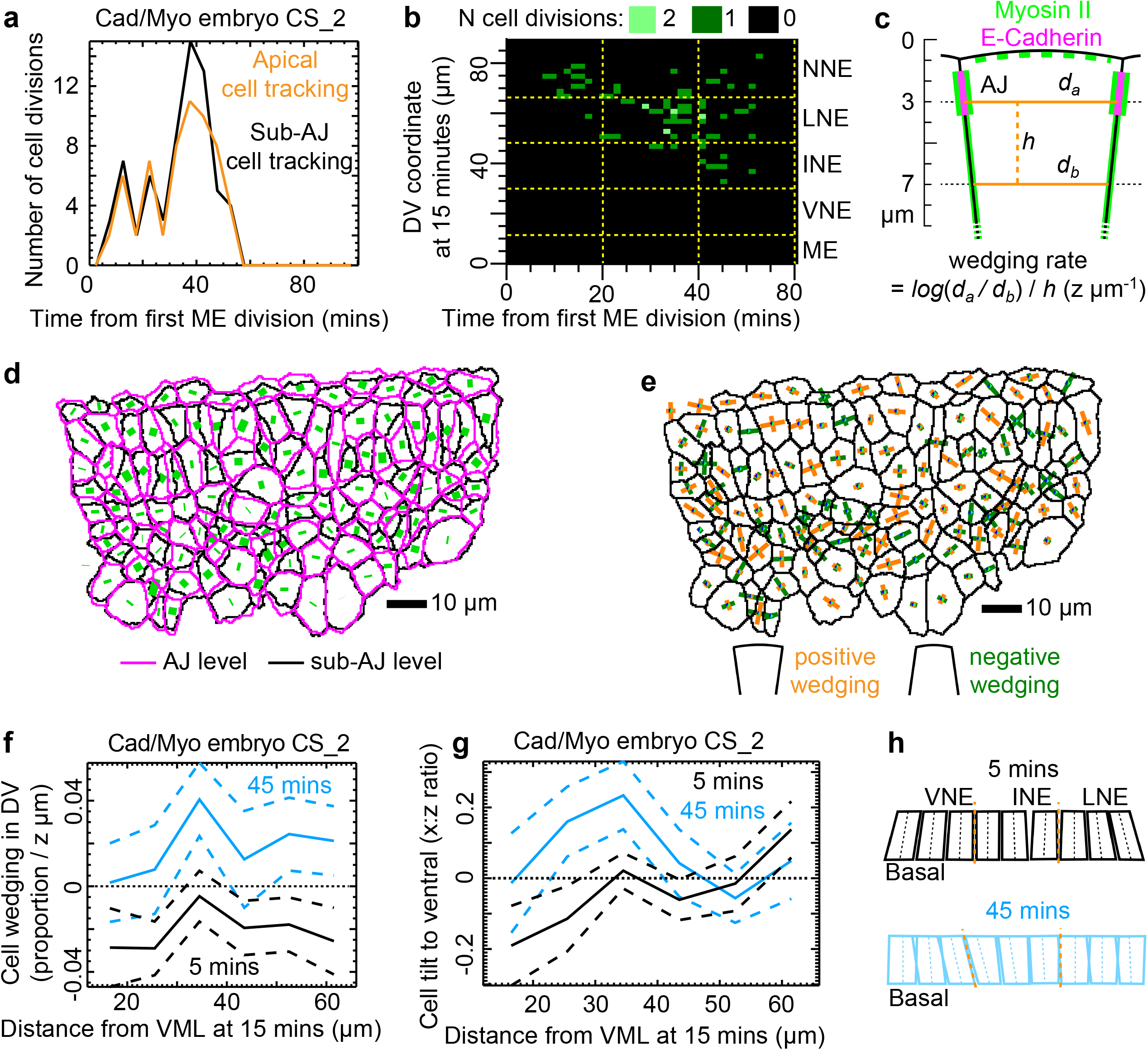
Cell tilt and wedging through tissue stress inversion. **a)** Frequency of cell divisions identified in cell tracking at AJ (−2.5 µm from epithelial surface) and sub-AJ (−6.5 µm) levels for embryo movie CS_2. A handful of extra divisions were found at the sub-AJ level, likely due to better manual tracking curation. **b)** Timing and DV location of divisions identified at the sub-AJ level (see Fig. 2Sh for AJ divisions for 5 Cad/Myo embryos). **c)** Schematic for how a wedging rate in depth is calculated from cell widths at AJ and sub-AJ depths. **d)** Overlay of cell segmentation of AJ (magenta) and sub-AJ (black) tissue layers for an example movie frame of embryo CS_2. Green lines connect cell centroids between levels. VML is at the top, with a line of ME cells then NE below. Anterior left. **e)** Principal axes of rates of cell wedging for cells in the tissue region shown in (d). **f)** Cell wedging at 5 and 45 mins, during and after the tissue stress reversal. During the stress reversal, cells are consistently more negatively wedged, irrespective of location along DV. **g)** Average cell tilt across the tissue DV axis for cells during the tissue stress reversal (5 mins) and afterwards (45 mins). Tilt towards ventral is positive. During the stress reversal, cells are tilted towards the middle of the tissue (negative tilt near the VML, positive near the NNE). **h)** Approximate summary cartoons of the data in (f) and (g). Side-views of the epithelium from ventral to dorsal (left to right) show approximate wedging and tilting patterns during the stress inversion (−5 mins) and after DV stress has been fully re-established (45 mins).

**Figure 5 Supplement.**
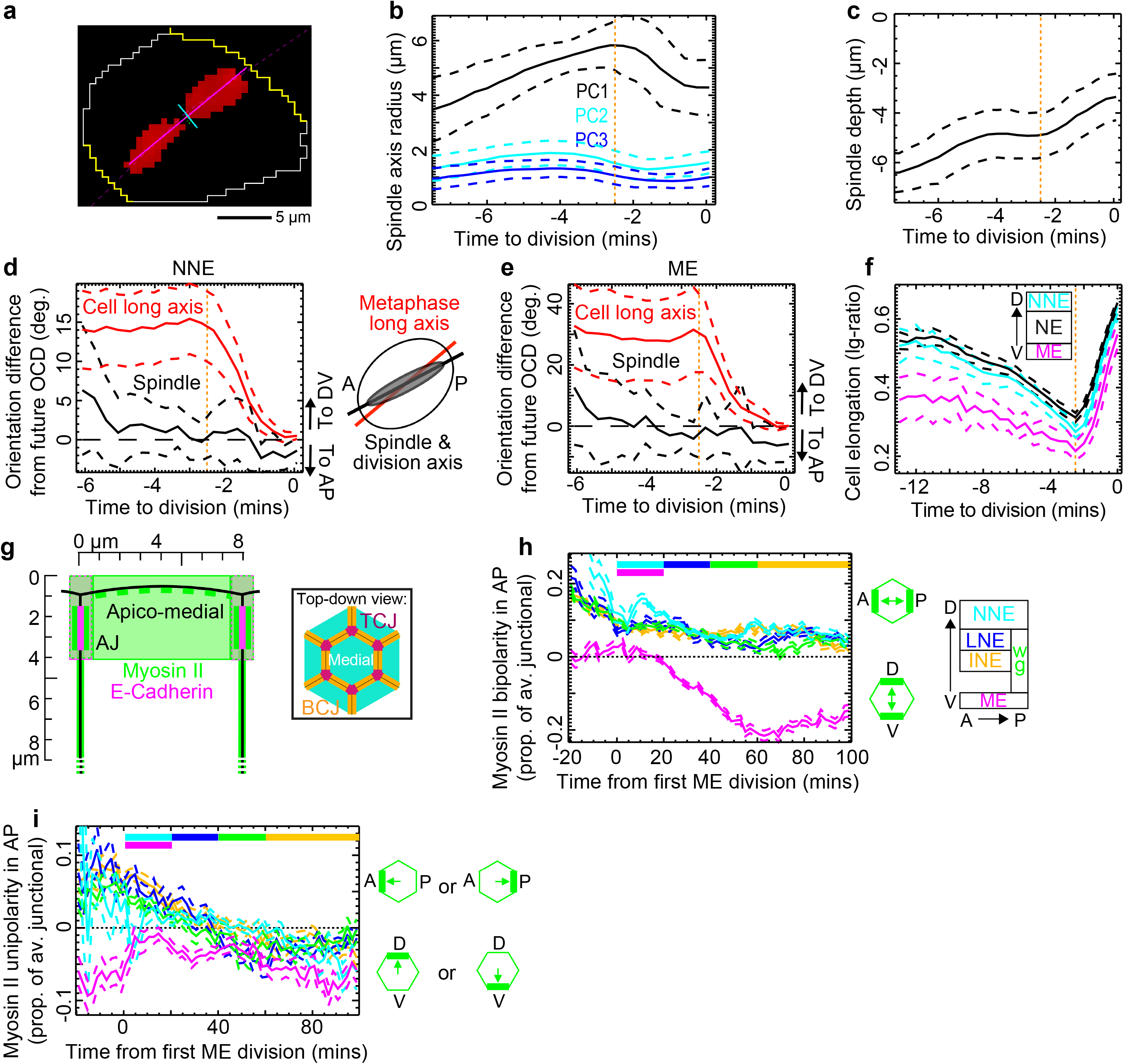
An AP-oriented cue attracts the mitotic spindle independently of mechanics. **a)** Example segmented cell outline (yellow) in late metaphase with pixels above threshold (red) identified as spindle pixels. **b)** 3D principal component analysis (PCA) of thresholded pixel volumes reveal elongated shapes with one dominant axis peaking in length at the start of anaphase (−2.5 mins). **c)** Average depth of the spindle centre for the same data as in (b). **d-e)** Angular difference, to AP or DV, between both the cell long axis and spindle and the OCD for **d)** NNE and **e)** ME cells. **f)** Average cell elongation for NE, ME and NNE cells during mitosis. **g)** Schematic of the apical depth range used to quantify Myosin II at BCJs, TCJs, and apico-medially (see inset and Methods). **h-i)** Myosin II **h)** bipolarity and **i)** unipolarity over developmental time by mitotic domain, projected onto AP axis. In all data panels, averages are plotted +/−95% Cis (dashed lines). Vertical orange dotted lines mark the metaphase to anaphase transition (−2.5 mins). Data pooled from 4 Cad/MT embryos in **b-e,** from 5 Cad/Myo and 4 Cad/MT embryos in **f** and from 5 Cad/Myo embryos in **h-i**.

**Figure 6 Supplement.**
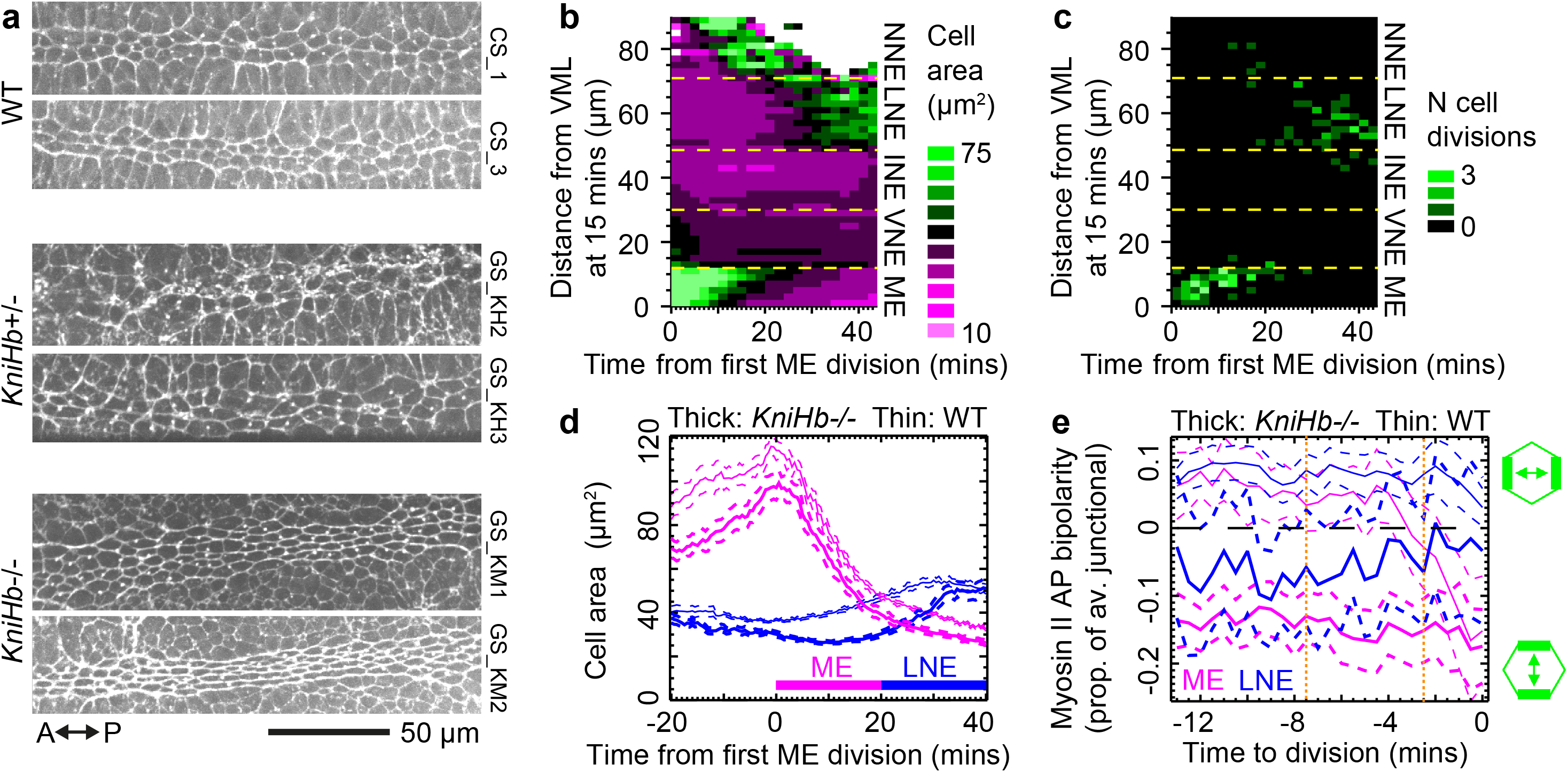
Anaphase long cell axis rotation towards AP is lost in an AP-patterning mutant. **a)** Stills from E-Cadherin channel from two further example embryos per genotype (see Fig. 6a) showing the width of the ME after division. **b-e)** Data pooled from three *KniHb-/-* embryos. **b)** Patterns of apical cell area over developmental time and across DV (compare to Fig. 2a and Fig. 2Sd). **c)** Patterns of cytokinesis events in ME and LNE domains over developmental time and across DV (compare with Fig. 2Sh, i). **d)** Average apical cell areas per mitotic domain over developmental time (see Fig. 6d). **e)** Average Myosin II bipolarity through mitosis by mitotic domain. *KniHb-/-* embryo data is negative on average, indicating DV-oriented bipolarity, compared to AP-oriented planar polarity in wild-type. Orange dotted lines at start of metaphase and anaphase.

**Figure 7 Supplement:**
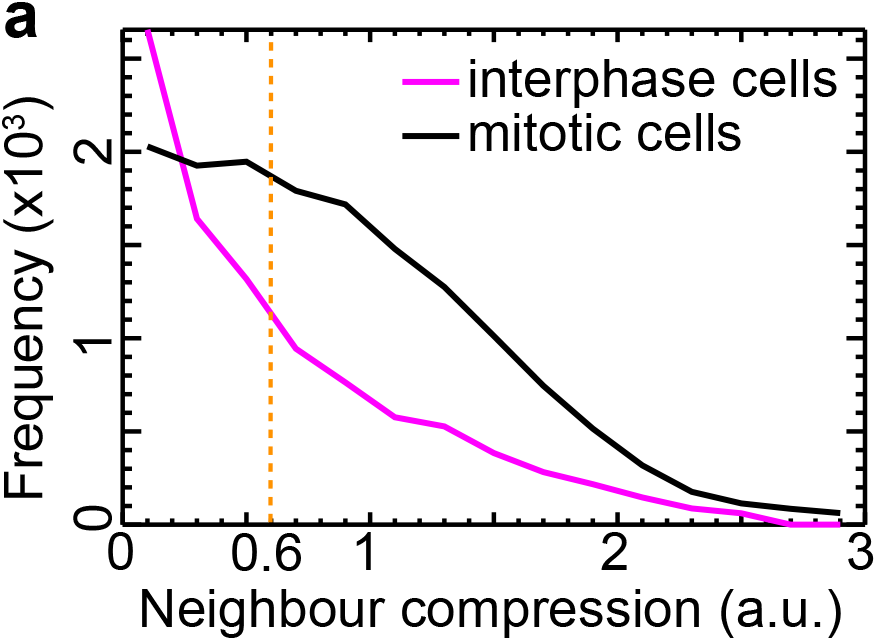
Compression from neighbouring dividing cells during metaphase re-orients divisions. **a)** Frequency of the strength of neighbour compression for all cells with at least one neighbour in mitosis. Because divisions occur in quasi-synchronous nests, neighbour compression is more frequent in mitotic than interphase cells. Orange dotted line, threshold above which neighbour compression re-orients focal cell long axes (see Fig. 7f).

**Supplementary Movie 1: Classifying cell division events in raw movies.** Top left; E-Cadherin (magenta) and Myosin II (green) fluorescence confocal movie channels. Top right; best-fit ellipses drawn on top of segmented and tracked cell shapes. Lengths of principal axes are colour-coded blue (1 µm) to green (5.5 µm) to red (10 µm). Bottom left; dumbbell pinch measure for identifying cytokinesis, increasing from zero (blue) to 0.2 (red). Bottom right; final classification of cytokinesis events. Cells are colour-coded by time to cytokinesis from blue (<= −10 mins) to red (>= 0 mins). Images and data from Cad/Myo embryo CS_4.

**Supplementary Movie 2: Classified cell division events.** All 172 cell divisions classified in Cad/Myo movie CS_4 are portrayed in 30 s frame intervals from −11 mins to 0 mins, the frame before the mother cell splits into two daughters. The mother cell centroid is set to the centre of each square neighbourhood, which has been rotated so that the orientation of division is horizontal (as in Fig. 1e). Magenta is Shotgun-GFP (E-Cadherin) and green is Sqh-mCherry (Myosin II). Cytokinesis rings can be seen appearing as vertical green lines in the centre of dividing cells at around −2.5 mins.

**Supplementary Movie 3: Embryo AP and DV coordinates.** Example embryo axial coordinates for movie CS_4. Left panel; colour-coded DV distance from ventral mid-line at 15 mins, carried forwards and backwards by cells as a co-moving coordinate frame. Blue is 0 µm, red is 84 µm from VML. Right panel; within-parasegment normalised AP coordinates. Blue is 0 at anterior PSB, red is 1 at posterior PSB. For a cell to be allocated a within-parasegment coordinate, the locations of PSBs to anterior and posterior of the cell must be recorded. The appearance and disappearance of blocks of cells in this movie panel reflects the gain or loss of information on one or both of the local PSBs.

**Supplementary Movie 4: Average Cad/Myo mitosis behaviour.** Average cell fluorescence through mitosis from −13 mins to cytokinesis (0 mins), at 30 s frame intervals. Left panel is at depth 0.5 µm below the surface of the epithelium with subsequent panels stepping 2 µm deeper to 10.5 µm in right panel. Magenta is E-Cadherin, green is Myosin II. Scale bar as in Fig. 3c.

**Supplementary Movie 5: Average Cad/MT mitosis behaviour.** Average cell fluorescence through mitosis from −13 mins to cytokinesis (0 mins), at 20 s frame intervals. Left panel is at depth 0.5 µm below the surface of the epithelium with subsequent panels stepping 2 µm deeper to 10.5 µm in right panel. Magenta is E-Cadherin, green is MTs. Scale bar as in Fig. 3d.

**Supplementary Movie 6: Jupiter spindle tracking.** 3D ellipsoid spindle shapes were extracted from volume images of intra-cellular (cytoplasmic) Jupiter fluorescence. Principal axes of spindle shape are drawn on each cell’s centroid, longest first in the colour sequence green, red and blue. Tracked cell shapes are in pink. Embryo move CJ_4 is shown.

